# The gut microbiome shapes pharmacology and treatment outcomes for a key anti-inflammatory therapy

**DOI:** 10.1101/2025.06.26.661733

**Authors:** Vanya Sofia Villa Soto, Alexandra L. Degraeve, Chloe M. Heath, Diego A. Orellana, Erin R. Reilly, Mohana Mukherjee, Jacob G. Brockert, Darren S. Dumlao, Rebecca B. Blank, Noah Perlmutter, Steven Yu, Judith Ashouri, Jose U. Scher, Andrew D. Patterson, Peter J. Turnbaugh, Renuka R. Nayak

**Affiliations:** Department of Medicine, Division of Rheumatology, University of California, San Francisco, San Francisco, CA, USA; Department of Microbiology, Pennsylvania State University, University Park, PA, USA; Department of Microbiology & Immunology, University of California, San Francisco, San Francisco, CA, USA; Chan-Zuckerberg Biohub-San Francisco, San Francisco, CA, USA; Department of Medicine, Division of Rheumatology, New York University, New York, NY, USA

## Abstract

The human gut microbiome encodes a formidable metabolic repertoire that harvests nutrients from the diet, but these same pathways may also metabolize medications. Indeed, large screens have revealed extensive microbial metabolism of drugs *in vitro*, but the pharmacologic and clinical repercussions of microbiota-mediated metabolism *in vivo* remain to be discerned. As a proof-of-concept, we investigate how human gut microbes contribute to *in vivo* pharmacology and efficacy of a key anti-inflammatory drug, methotrexate (MTX). Specifically, we demonstrate that the gut microbiome shapes drug pharmacology *in vivo* in mice, both by directly metabolizing the drug and by inducing host pathways that promote drug metabolism. Moreover, interindividual variation in the human gut microbiome contributes to variation in pharmacokinetic (PK) profiles. When we quantified metabolites produced by microbes, we unexpectedly identified novel MTX metabolites, one of which, p-methylaminobenzoyl-L-glutamic acid (pMABG), was a major byproduct of microbial metabolism both *in vitro* and *in vivo*. Further, we find that a large proportion of patient-associated microbes are capable of metabolizing MTX. Finally, we show that microbial metabolism of MTX is linked to PK profiles and disease outcomes in a mouse model of inflammatory arthritis. Taken together, these findings provide evidence that the human gut microbiome causally contributes to drug pharmacology *in vivo* for a key anti-inflammatory drug through known and novel mechanisms. Our studies provide a framework for elucidating the clinical relevance of drug microbial metabolism in the context of treatment response. These results are a first step towards understanding and manipulating the human gut microbiome in the treatment of autoimmunity and the advancement of precision medicine for millions of patients taking MTX for immune or inflammatory conditions.

**Highlights:**

- The gut microbiome impacts methotrexate (MTX) pharmacology in mice
- The human gut microbiome contributes to interindividual variation in MTX pharmacology
- Human gut microbes produce novel MTX metabolites, pMABG and 6-MPDA
- Microbial metabolism of MTX is linked to treatment outcomes

## Introduction

The human gut microbiome encodes for millions of genes, many of which provide critical benefits for the host, but these microbial pathways may lead to unintended consequences in the setting of drug therapy. For example, microbes facilitate nutrient extraction^1^, but they also metabolize therapeutic drugs^2^. Indeed, this extensive metabolic capacity of microbes has been demonstrated in recent large-scale *in vitro* drug screens: >400 therapeutic drugs were metabolized by individual microbial isolates as well as by complex bacterial communities from healthy individuals *ex vivo*^3,4^. However, the *in vivo* repercussions (i.e., therapeutic efficacy or toxicity) of this metabolism on the host remain largely unexplored. Thus, translational studies are needed to elucidate how the microbiome contributes to treatment outcomes in the host. They are essential for developing microbiome-based interventions to transform treatment^5,6^.

Toward this goal, we chose to investigate an anchor therapy in the treatment of autoimmune and inflammatory disease: methotrexate (MTX). From a microbiome perspective, MTX is an ideal drug to study because it is an antimetabolite: it structurally mimics an essential vitamin, folate, that is required by nearly all forms of life^7,8^. Microbes encode enzymes to synthesize^99^ and break down folate (unlike mice or humans)^10,11^. Importantly, these same folate metabolism enzymes act on MTX^12^. Thus, compelling *in vitro* evidence supports the hypothesis that the microbiome shapes MTX pharmacokinetics (PK) and impacts treatment outcomes.

From a clinical perspective, MTX is widely used to treat millions of patients suffering from autoimmunity, cancer, and other autoimmune diseases (e.g., dermatomyositis, lupus, psoriasis and psoriatic arthritis)^13^. Hence, it is designated by the World Health Organization (WHO) as an essential medicine. Currently, oral MTX is first-line treatment for rheumatoid arthritis (RA)^14^ and confers multiple unique therapeutic benefits to patients^15^. For these reasons, treatment guidelines recommend maximally up-titrating oral MTX or, if that fails, using subcutaneous MTX, which is more bioavailable^14^. Thus, MTX is a central therapeutic in the treatment of autoimmunity, and maximizing its efficacy is important for multiple reasons.

Patients, however, show extensive interindividual variation in MTX response, and the underlying molecular mechanisms remain incompletely understood despite decades of investigation^16–18^. Host and environmental factors have been explored with the goal of identifying modifiable factors to improve response^16^. While genetic factors and smoking have been found to contribute to MTX responsiveness, other factors likely remain unidentified^16^.

Our group focuses on the impact of the microbiome on the treatment of autoimmunity. We recently showed that among RA patients, pre-treatment microbiomes differ between MTX-responders (MTX-R) versus MTX-non-responders (MTX-NR)^19^. In *ex vivo* studies, pre-treatment RA microbiota variably depleted MTX, and depletion was negatively associated with future MTX response^19^. We identified multiple human gut bacterial strains that directly deplete MTX *in vitro*^20^. Together, these findings show that human gut microbes directly metabolize MTX and that this metabolism is associated with clinical response in patients. However, demonstration that microbial metabolism causally impacts MTX PK and shapes treatment outcomes remains to be determined. Such knowledge is required to translate fundamental basic science findings and pave the way for rationally manipulating the gut microbiome to maximize efficacy of this central therapeutic.

The role of the gut microbiome in MTX pharmacology has been previously explored in animal models^21^, but whether human gut microbiota metabolize MTX has been debated^22,23^ or metabolism was thought to be minimal^24^. These early studies were conducted when MTX was given intravenously, but currently it is administered orally, theoretically allowing for more interactions between MTX and the gut microbiome. Furthermore, early studies were done before the development of highly sensitive analytical chemistry platforms, which could distinguish between MTX and its metabolites. Hence, for historical reasons, previous studies may have underestimated MTX-microbiota interactions and the ability of the microbiota to shape drug pharmacology and drug response.

Given these clinical associations and supportive *ex vivo* and *in vitro* findings, here we test the causal contributions of gut microbial metabolism to MTX response *in vivo*. Specifically, we test the extent to which the human gut microbiome contributes to MTX PK, identify the spectrum of human microbes that metabolize MTX and the metabolites they produce, and examine the impacts of microbial metabolism on treatment outcomes. Distinct from prior studies^21,25^ largely examining fecal and urinary drug levels and murine microbiota, we quantify levels of MTX and its metabolites in the circulation of gnotobiotic mice colonized with human gut microbes. We find that the microbiome causally contributes to MTX PK and that this metabolism is linked to treatment outcomes in mice. These findings raise the possibility that microbial metabolism can be modulated to impact MTX outcomes in the host. These results are a first step towards manipulating the human gut microbiome in order to advance precision medicine for millions of patients globally taking MTX for autoimmunity.

## Results

### The gut microbiome impacts MTX pharmacology *in vivo*

We first sought to determine whether the gut microbiome contributes to MTX PK *in vivo*. To test this, we compared plasma MTX levels in germ-free (GF, N=6) vs. specific-pathogen-free (SPF, N=8) C57BL/6J mice given MTX 50 mg/kg oral dose (**Supplementary Figure 1A**). We also quantified levels of 3 previously reported MTX metabolites: deoxyaminopteroic acid (DAMPA), 7-hydroxy-methotrexate (7-OH-MTX), and polyglutamated MTX (MTXPG_n_) (**Figure 1A**)^15^. Of these previously reported MTX metabolites, DAMPA is the only one that is exclusively produced by microbes^26^. 7-OH-MTX is produced by the host liver enzyme aldehyde oxidase (AO)^27,28^; to the best of our knowledge, this is not produced by microbes. Polyglutamated MTX metabolites, ranging from 2 to 7 additional glutamates, are primarily produced once MTX is absorbed into host cells and can accumulate intracellularly over several days to months^29^. We collected tail vein blood at 0 hours (prior to administration) and multiple time points after dosing and profiled plasma concentrations. From these PK curves, Area Under the Curve (AUC) and peak concentrations were calculated for each mouse.

**Figure 1.**
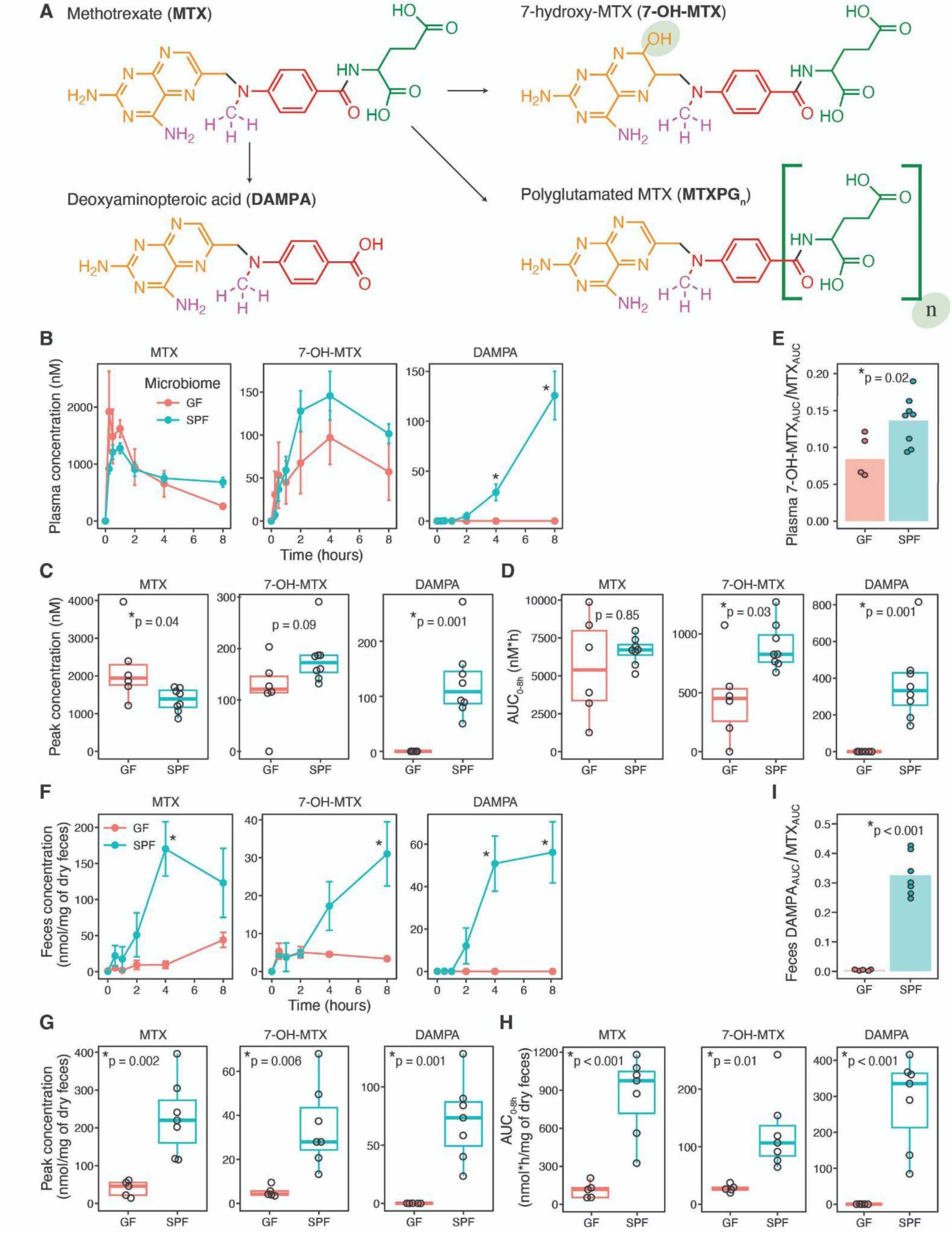
The gut microbiome impacts MTX PK *in vivo*. (**A**) Structures of methotrexate (MTX) and its known metabolites, 7-hydroxy-MTX (7-OH-MTX), deoxyaminopteroic acid (DAMPA), and polyglutamated MTX (MTXPG_n_). (**B**) Plasma concentrations of MTX and its known metabolites were quantified by LC-MS in germ-free (GF, N=6) or colonized (SPF, N=8) C57BL/6J female mice after a one-time oral administration of MTX 50 mg/kg. Multiple plasma samples were taken from each mouse over an 8 h time course. (**C**) Peak plasma concentration, and (**D**) Area Under the Curve (AUC) were compared between GF and SPF mice. (**E**) Ratio of AUC of 7-OH-MTX to MTX in the plasma over 8 h. (**F**) Stool concentrations of MTX and its metabolites were quantified at multiple time points. (**G**) Peak stool concentrations, (**H**) AUC and (**I**) ratio of AUC of DAMPA to MTX are shown. Results from Student’s t-test (**B-I**) are reported. Error bars are SEM.

Peak plasma concentrations of MTX were significantly higher in GF mice relative to SPF mice (**Figure 1B-C**), suggesting that the microbiome reduces MTX absorption from the gut. Peak plasma DAMPA concentrations were significantly higher in SPF mice and notably absent in GF mice (**Figure 1C**), consistent with previous reports indicating that DAMPA is exclusively microbially produced^30^. This finding also shows that while DAMPA is produced in the gut, it can enter host circulation. The mean AUC of 7-OH-MTX was higher in SPF mice compared to GF mice (**Figure 1D**). To discern whether this might be due to increased AO enzymatic activity in SPF mice, we compared the product-substrate ratio between 7-OH-MTX and MTX. The ratio of 7-OH-MTX to MTX (7-OH-MTX_AUC_/MTX_AUC_) was significantly higher in SPF mice (**Figure 1E**), suggesting that the microbiome increases MTX liver metabolism in the host. Finally, no significant differences were observed for polyglutamated MTX (**Supplementary Figure 1B**), which was either absent or present at very low levels, likely because multiple weeks are required for accumulation of this metabolite^29^.

We next quantified fecal levels of MTX and its metabolites. MTX levels were significantly higher in SPF mice compared to GF mice (**Figure 1F-H**). Consistent with it being a microbiota-produced metabolite, DAMPA was only seen in the feces of SPF mice (**Figure 1F-H**) and was present at high levels, suggesting incomplete absorption of DAMPA. Similar to plasma levels, fecal 7-OH-MTX concentrations were higher in SPF in mice than in GF mice (**Figure 1F-H**). These findings in the stool support the hypothesis that the gut microbiota impacts drug metabolism to shape MTX pharmacology.

Taken together, these results suggest that the mouse gut microbiome contributes to MTX PK *in vivo* through microbiota- and host-mediated mechanisms. We show for the first time that DAMPA, which is exclusively produced by gut microbes, enters into host circulation. We also show for the first time that levels of 7-OH-MTX, which is made by the liver and excreted into the gut, is impacted by the microbiota. Thus, the microbiome contributes to MTX pharmacology directly, through microbial metabolism, and indirectly, through mechanisms of absorption, metabolism, and/or enterohepatic circulation.

### Parenterally administered MTX undergoes microbial metabolism

Next, we considered whether the route of administration might mitigate microbiota-mediated metabolism of MTX. Because MTX can be administered orally or subcutaneously in patients with autoimmune disease, we sought to determine whether gut microbial metabolism of MTX could be bypassed by administering MTX through non-oral routes. We treated SPF C57BL/6J mice with MTX 50 mg/kg either via oral gavage (PO) or by intraperitoneal injection (IP) (N=3 / route of administration). Peak plasma MTX levels and AUC were significantly higher in mice given IP MTX compared to PO (**Supplementary Figure 2A-C**), a finding that is consistent with prior reports that parenteral MTX has higher bioavailability^31^. Similarly, peak 7-OH-MTX levels and AUC were elevated in mice given IP dosing (**Supplementary Figure 2A-C**); the ratio of 7-OH-MTX to MTX did not differ by route (**Supplementary Figure 2D**). Plasma concentrations of DAMPA were detected in both groups and did not significantly differ (**Supplementary Figure 2A-C**). These findings suggest injection-dosing of MTX does not bypass gut microbial metabolism, likely due to enterohepatic circulation of the parent compound^32^.

### Interindividual variation in human gut microbiota contributes to MTX PK *in vivo*

Having established a role for the murine microbiome in MTX PK, we next sought to determine the impact of the human gut microbiome on MTX PK and to ascertain the clinical relevance of our findings. Specifically, we asked whether natural interindividual variation in human gut microbial communities of patients contributes to MTX PK *in vivo*. We focused on RA patients because MTX is first-line therapy for this common autoimmune disease affecting 1% of the worldwide population^1414^. We colonized GF C57BL/6J mice with fecal communities from two different treatment-naïve RA patients (Patient 170 vs. Patient 317, N=11 mice per donor colonization group) and treated with oral MTX 50 mg/kg. Mice colonized with Patient 170 microbiota had significantly higher peak plasma concentrations of MTX than mice colonized with microbiota from Patient 317 (**Figure 2A-B**). We were unable to detect significant differences between 7-OH-MTX levels or 7-OH-MTX/MTX ratios between colonization states, suggesting similar levels of first-pass liver metabolism (**Figure 2B-E**). While there was a trend toward lower plasma levels of DAMPA in mice colonized with Patient 170 microbiota (consistent with the higher MTX levels in this colonization group), we were unable to demonstrate that this difference was statistically significant (**Figure 2B-D**). These findings show that interindividual variation in RA patient microbiota contributes to differences in MTX PK *in vivo*.

**Figure 2.**
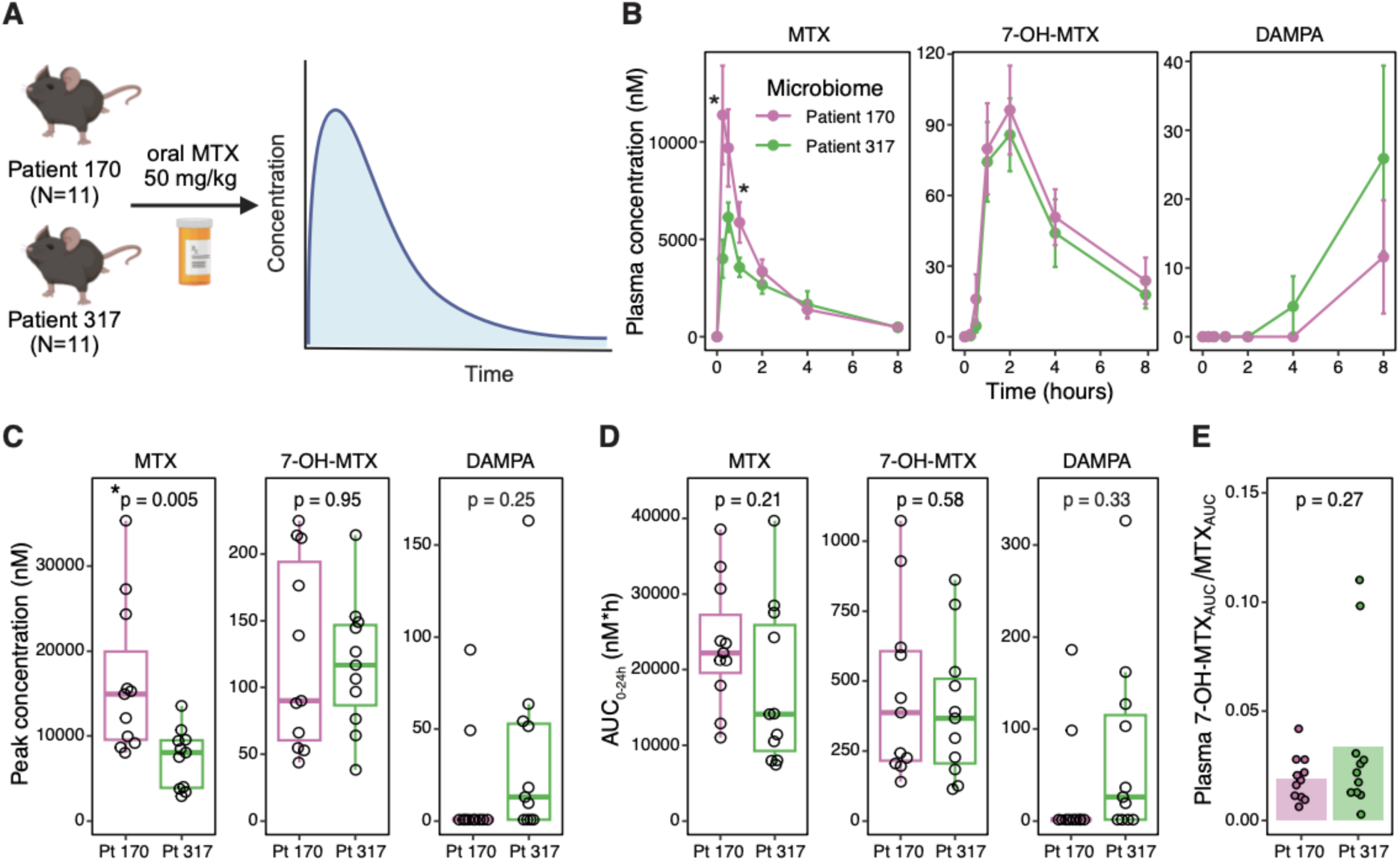
Interindividual variation of the human gut microbiome contributes to MTX PK *in vivo*. (**A**) Experimental design of PK study in GF C57BL/6J male mice colonized with donor microbiota from two RA patients (Patient 170 and Patient 317; N=11 per colonization group) given a one-time oral administration of MTX 50 mg/kg. (**B**) Plasma concentration of MTX and its known metabolites were quantified by LC-MS. Multiple plasma samples were taken from each mouse over a 24 h time course (first 8 h depicted for clarity). (**C**) Peak concentration and (**D**) AUC were compared between mice colonized with gut microbiomes from Patient 170 or Patient 317. (**E**) Ratio of AUC of 7-OH-MTX to MTX in the plasma over 24 h. Results from Student’s t-test are reported. Error bars are SEM.

### Identification of novel MTX metabolites produced directly by human gut microbes

We were perplexed that despite finding differences in plasma MTX concentrations in mice colonized with different microbiota, we did not find concomitant statistically significant differences in either DAMPA or 7-OH-MTX (**Figure 2**). This led us to ask whether MTX may be metabolized into novel compounds that were not captured/quantified in our studies. In prior studies, we found that type strains of *Clostridium asparagiforme*, *Clostridium innocuum*, and *Clostridium symbiosum* depleted MTX from the culture medium as assessed by high-performance liquid chromatography (HPLC)^20^. But when we profiled these cultures via liquid chromatography-mass spectrometry (LC-MS), which is able to disambiguate between MTX metabolites, we were surprised to find that while MTX was depleted by all three strains, only *C. symbiosum* produced DAMPA, and in relatively small amounts (**Figure 3A**). This led to two non-mutually exclusive hypotheses: (1) DAMPA is further metabolized by *C. asparagiforme* and *C. innocuum*, which is why we do not detect it, or (2) these two species metabolize MTX into metabolites other than DAMPA.

**Figure 3.**
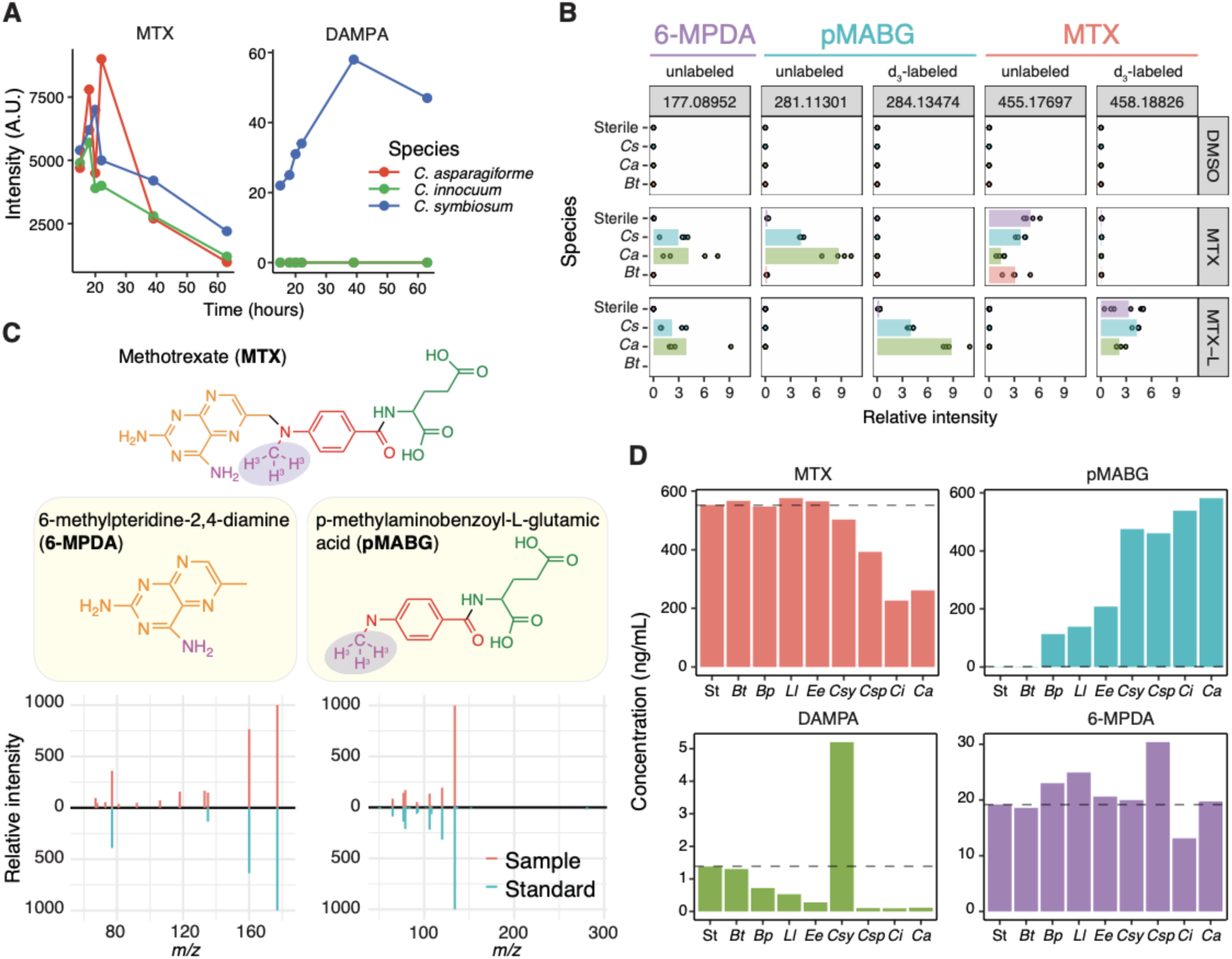
Human gut microbes produce novel MTX metabolites. (**A**) MTX and DAMPA peak intensities (peak analyte areas) in the supernatant of three type strains incubated with MTX over 63 h. (**B**) Stable isotope profiling of three type strains incubated with MTX or d_3_-labeled MTX (“MTX-L”) for 72 h (N=4 replicates per species per compound). Feature peak intensities that correspond to labeled and unlabeled metabolites are depicted in the bar plot. Intensity is normalized to the mean (average of all samples) and relative intensity is reported. The mass-to-charge (*m/z*) value of each peak is listed on top, and the treatments are denoted on the right-hand side. (**C**) Chemical structures of the 2 novel microbiota-derived metabolites of MTX identified by stable isotope labeling experiments (upper panel). The purple shading indicates the location of the deuterium label. Novel metabolites were confirmed using authentic standards: mirror plots comparing the sample (red) with the authentic standard (blue) are shown (lower panel). (**D**) Quantification of MTX and its microbial metabolites from the supernatant of multiple human gut bacterial isolates incubated with MTX (100 ug/mL) for 72 h *in vitro*. Dashed grey lines represent the amount of compound in sterile control wells. St, sterile; *Bt*, *Bacteroides thetaiotaomicron*; *Bp*, *Blautia producta*; *Ll*, *Lactonifactor longoviformis*; *Ee*, *Eubacterium eligens*; *Cs*, *Clostridium symbiosum*; *Csy*, *Clostridium symbiosum*; *Csp*, *Clostridium sporogenes*; *Ci*, *Clostridium innocuum*; *Ca*, *Clostridium asparagiforme*.

To determine the metabolic fate of MTX, we performed stable isotope labeling experiments followed by global mass spectrometry profiling. Specifically, we incubated unlabeled and d_3_-labeled MTX with 3 human gut bacterial isolates, 2 of which metabolize the drug (*C. asparagiforme* and *C. symbiosum*) and 1 which does not (*Bacteroides thetaiotaomicron*)^20^. After 72 hours of incubation, stable isotope profiling identified 2 novel MTX metabolites. While DAMPA was not detected in this isotope labeling study perhaps because it was not produced in sufficient amounts, we found evidence of a novel metabolite carrying the heavy isotope (**Figure 3B**): p-methylaminobenzoyl-L-glutamic acid (pMABG) with a mass-to-charge (*m/z*) value of 281 (unlabeled) or 284 (d_3_-labeled). Based on chemical logic, we surmised that a second unlabeled, novel compound should be produced upon generation of pMABG: 6-methylpteridine-2,4-diamine (6-MPDA) with a *m/z* value of 177. We obtained authentic standards of these compounds and confirmed that the fragmentation pattern of our novel metabolites matched those of the authentic standards (**Figure 3C**). We next asked which of these novel metabolites are produced by other species previously shown to metabolize MTX^20^, and found that many of these species produced predominantly pMABG rather than DAMPA (**Figure 3D**), suggesting that pMABG is a major microbial metabolite of MTX.

### Novel MTX metabolites are produced by complex human microbial communities and detected *in vivo* in mice

We next sought to determine whether these novel metabolites are produced by complex microbial communities from RA patients. We incubated fecal slurries from 17 treatment-naïve patients with vehicle control versus MTX 100 Ιg/mL for 24 hours and profiled the supernatant by using a newly developed and validated targeted LC-MS method. We found that RA microbiota produced significant amounts of DAMPA and pMABG, with pMABG being the major metabolite (**Figure 4A**).

**Figure 4.**
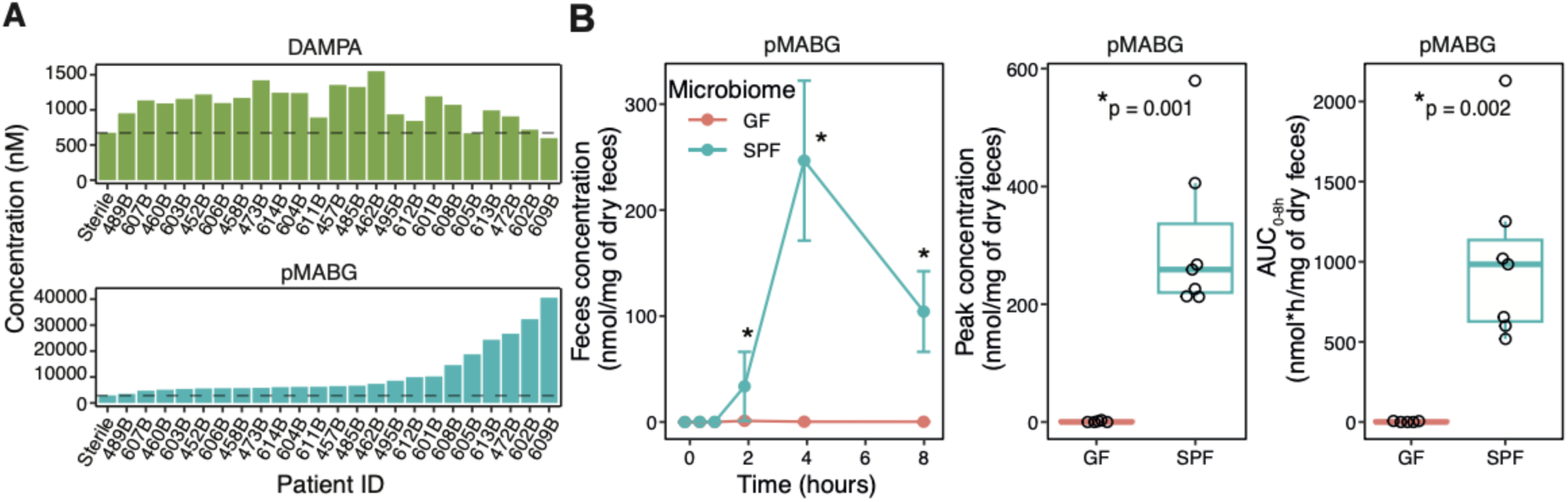
Novel metabolites are produced by human gut microbiota *ex vivo* and by murine microbiota *in vivo*. (**A**) Microbial communities from treatment-naïve RA patients produce novel MTX metabolites *ex vivo* to variable extents. Fecal slurries were prepared in pre-reduced saline and incubated with 100 mg/mL of MTX or vehicle control for 24 h in an anaerobic chamber. “Sterile” refers to sterile media with MTX. (**B**) As in Figure 1, fecal samples from GF (N=6) or colonized (SPF, N=8) C57BL/6J female mice were profiled on LCMS to quantify the novel metabolites (6-MPDA unable to be quantified). Results from Student’s t-test are reported. Error bars are SEM.

To determine whether pMABG is produced by the host or exclusively produced by microbiota, we returned to the fecal samples harvested from our initial gnotobiotic mouse experiment and found high levels of pMABG in the stool of SPF mice but not GF mice (**Figure 4B**). These findings suggest that pMABG is exclusively microbially produced.

Taken together, these results show that human gut microbes metabolize MTX into at least 3 compounds: DAMPA, pMABG, and 6-MPDA. DAMPA has not been previously shown to be directly produced by human gut bacteria (as we have shown here with *C. symbiosum in vitro* and *in vivo* in mice, **Figure 2**), and the other 2 have never been reported to be MTX metabolites before. Thus, these studies have uncovered novel metabolites of a widely used antirheumatic drug that has been used for over 4 decades, demonstrating that it is only with recent advances in anaerobic microbiology and analytical chemistry that we are now able to identify novel MTX metabolites.

### Identification of multiple novel MTX metabolizing bacteria from RA patients

We next sought to determine whether bacterial species harvested from RA patient microbiota could similarly metabolize MTX into known and novel metabolites. To build a culture collection of microbial strains from RA patients, we harvested bacterial isolates from 12 treatment-naïve RA patients. Patients provided stool samples prior to initiating treatment. We used rich media to maximize recovery of multiple species and strains as well as a spore-forming enrichment method^33^ to increase the odds of harvesting *Firmicutes*, which our prior studies suggested are enriched for metabolizing species^20^. Colonies with varying morphologies were identified and picked for further testing (**Supplementary Figure 3**).

We harvested a total of 360 bacterial isolates from RA patient microbial communities (median of 24 isolates per patient; range: 24 – 48). Of these, 306 (306/360, 85%) grew in liquid media and could be further studied; 54 (54/360, 15%) grew on the agar media but not in liquid media (**Supplementary Figure 4A**). Of the 306 bacterial isolates that grew in liquid media, 221 (221/306, 72**%**) grew in MTX 50 mg/mL and could be assessed for MTX metabolism capacity. The remaining 85 (85/306, 28%) likely did not tolerate the growth inhibitory effects of MTX 50 ug/mL.

Next, we sought to identify these strains. Of the 360 total isolated strains, we were able to identify bacterial species by full-length 16S sequencing for 241 isolates (241/360, 67%). These represented 36 unique species (**Supplementary Figure 4B**). From each patient, we identified 2-15 unique species (median = 7).

Next, we asked which of our harvested isolates metabolize MTX *in vitro*. Of the 221 isolates that grew in MTX, we identified 63 (63/221 = 29% of the isolates that grew in MTX or 63/360 = 18% of total isolates harvested) that could metabolize MTX, defined as strains showing at least a 60% depletion of MTX in the supernatant (**Figure 5A**). These represented at least 12 unique species as determined by full-length 16S sequencing (**Figure 5A**). Other than *Clostridium innocuum*^20^, none have previously been reported to metabolize MTX. Five unique species seemed to exhibit strain-level variation in MTX metabolism, in which multiple isolates of the same species variably showed 60% depletion. These included *Clostridium cibarium*, *Coprobacillus cateniformis*, *Romboutsia hominis*, *Enterococcus faecalis*, and *Intestinibacter bartletti*. These findings suggest that multiple patient-associated gut bacterial species can metabolize MTX.

**Figure 5.**
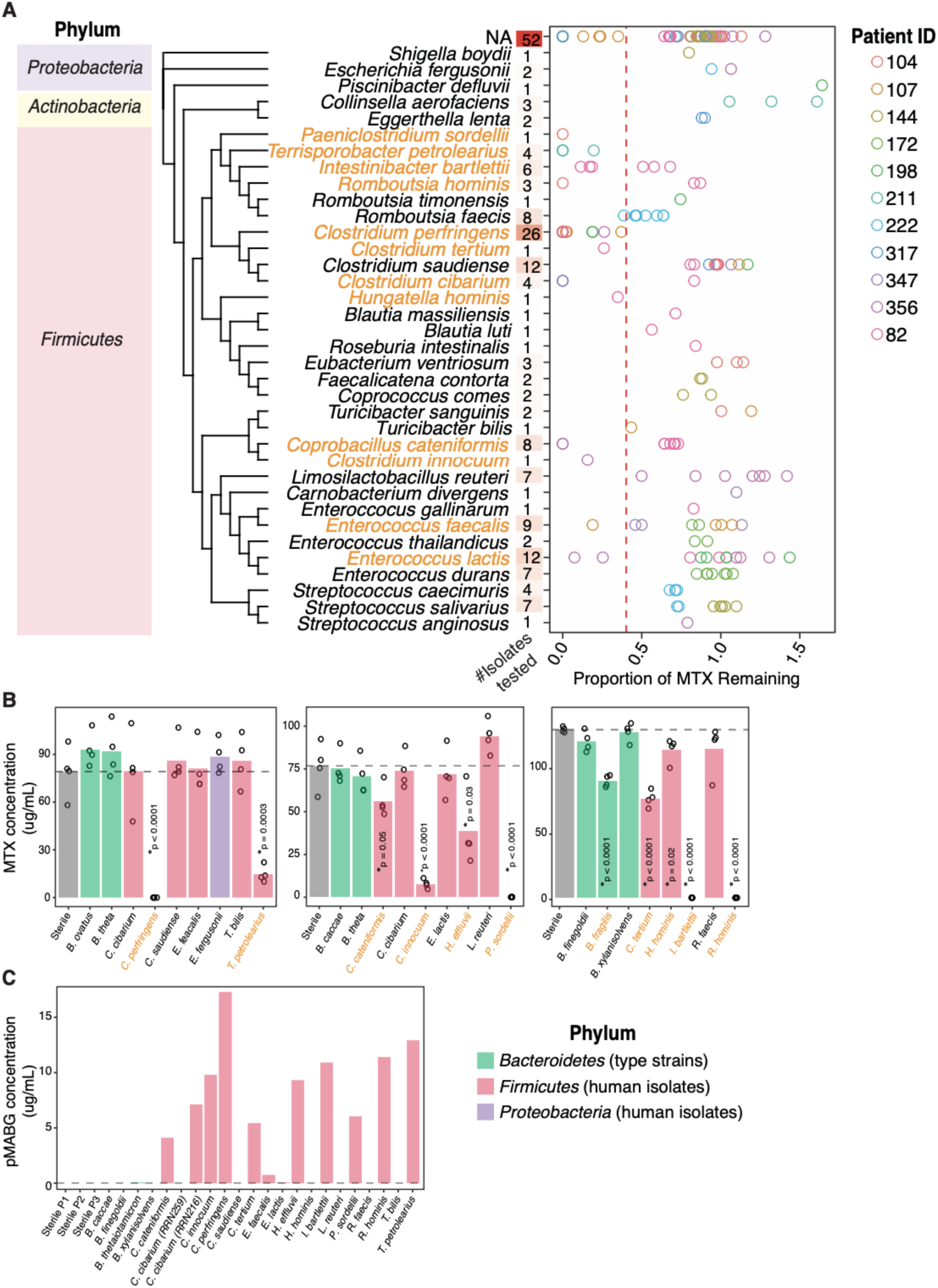
Multiple human gut bacterial strains isolated from treatment-naïve RA patients can metabolize MTX *in vitro*. (**A**) Results of our MTX metabolism screen showing the phylum and species identification (left), the number of strains per species tested (middle), and the proportion of MTX remaining in the supernatant relative to sterile controls (right). The RA patient donor of each strain is also shown (indicated by different color circles). Strains that depleted MTX by at least 60% (red dashed line) were scored as “metabolizers” and are highlighted in orange. (**B**) Validation of MTX depletion by each human isolate. Each isolate was incubated with MTX 100 ug/mL for 72 h (N=4 replicates per species) and MTX was quantified by HPLC. RA-associated species that significantly deplete MTX are highlighted in orange. (**C**) LC-MS quantification of pMABG in the same bacterial supernatant as in (**B**). For (**B**) and (**C**), the dashed line indicates abundance of the compound in sterile controls. The phylum is indicated by the color of the bar. Results of Student’s t-test are reported.

### A novel metabolite is a major product of microbial MTX metabolism

Next, we sought to identify the metabolites produced by each of these 12 unique species. For 10 of the 12 strains, we validated significant MTX depletion compared to sterile and negative controls as quantified by HPLC (see Methods)(**Supplementary Table 1, Figure 5B**); 1 showed impaired growth (*E. faecalis*, 70% compared to vehicle control) and so we may have been limited in our detection of MTX metabolism, and 1 (*Enterococcus lactis*) failed to validate. For all 10 validated strains, LC-MS revealed that pMABG was the dominant metabolite produced (**Figure 5C**). While *E. faecalis* did not show significant depletion of MTX by HPLC (**Figure 5B**), we did detect modest pMABG production (**Figure 5C**), suggesting that it is a MTX metabolizer. For all 11 of the metabolizing strains, we did not detect 7-OH-MTX or DAMPA production compared to sterile controls (**Supplementary Figure 4C**). This may be either because these are not produced or because they are further metabolized and shuttled into other anabolic pathways. Given that 6-MPDA is expected to be a byproduct of pMABG production, we were expecting to see elevated levels of 6-MPDA in metabolizers. However, we were not able to detect significant elevations of 6-MPDA compared to controls (**Supplementary Figure 4C**), suggesting either that our method for extracting and quantifying this small molecule needs refinement or that a compound other than 6-MPDA is produced upon pMABG generation (e.g., 6-MPDA bonded to a formyl or methyl group). While *Hungatella hominis* showed significant MTX depletion by HPLC compared to sterile controls, the extent of metabolism was modest (12% MTX depletion), and we were not able to detect any of the known or novel MTX metabolites by LC-MS. This might suggest that *H. hominis* makes metabolites that remain unknown or that the known metabolites are further metabolized into compounds that are not captured by our targeted method.

Taken together, we have identified at least 11 MTX metabolizing microbes harvested from RA patients (11/36, representing 31% of identifiable isolates) and found that the major metabolite produced is pMABG. These findings reveal that a significant percentage of strains isolated from RA patients have the capacity to metabolize MTX, with some strains exhibiting strain-level variation. These findings suggest that variation in gut microbiome composition and/or strain level variation could contribute to variation in MTX response via microbial drug metabolism.

### MTX response in an arthritis model is associated with the microbiome

Concurrent with our *in vivo* pharmacologic and *in vitro* metabolism studies above, we sought to develop an animal model system to test the impact of microbial metabolism on MTX efficacy *in vivo*. For this, we turned to the SKG mouse model of inflammatory arthritis, which recapitulates multiple features of human RA disease, including production of rheumatoid factor and CCP antibodies, development of chronic inflammatory arthritis of the small joints, elevated inflammatory markers, and female predominance^34–37^. Multiple previous studies have shown that gut microbes are important for the pathogenesis of autoimmunity in this model^34,35,38^, but none have explored its significance in therapeutic drug response.

Because prior studies showed that MTX alleviates SKG arthritis when given via the intraperitoneal route^39^, we first tested whether oral MTX similarly alleviated arthritis. In order to increase our sample size for this pilot experiment, we pooled SKG mice that were bred at two different facilities: Parnassus (“Parn.”) and Mount Zion (“Zion”). The mice from Zion were transferred to Parnassus, were cohoused with Parnassus-reared mice, and were permitted to acclimate to the Parnassus colony for 2 weeks (“Experiment 1”). Arthritis was induced in mice with a single IP injection of zymosan, and then mice were treated with oral MTX 10 mg/kg (N=8) or vehicle control (VEH, N=8) every other weekday for 3 weeks. The Clinical Arthritis Score (CAS) was significantly lower in mice treated with MTX compared to VEH mice (**Figure 6A-B**). These findings confirmed that oral MTX, similar to IP MTX^39^, alleviates arthritis in SKG mice.

**Figure 6.**
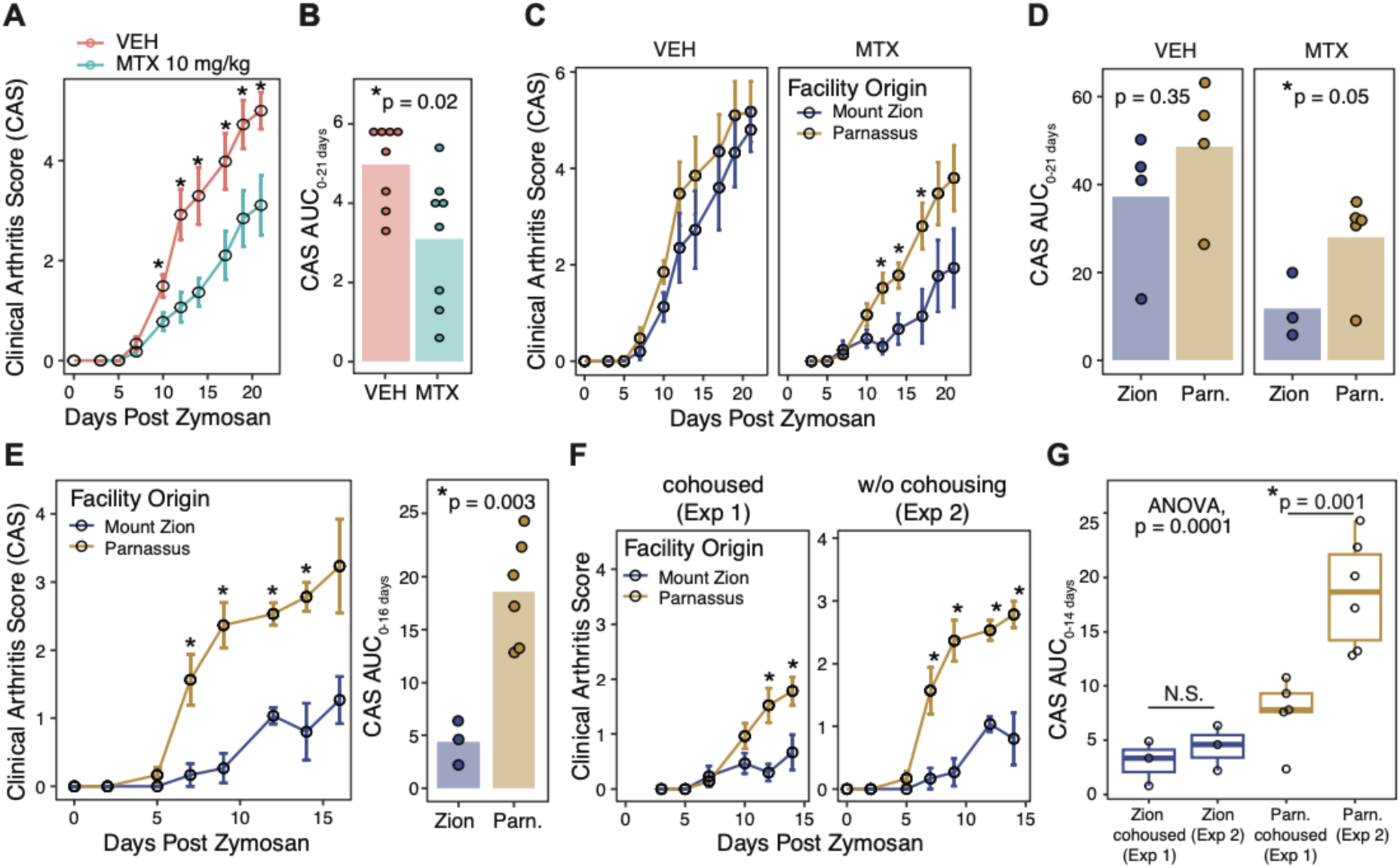
MTX response in a model of inflammatory arthritis is associated with the microbiome and is transferable. (**A**) Clinical arthritis scores (CAS) of female SKG mice treated with vehicle (VEH, N=8) or oral MTX 10 mg/kg (N=8) three times weekly. (**B**) AUC of CAS (from day 0 to day 21). (**C**) CAS over time and (**D**) AUC of CAS stratified by treatment (VEH vs. MTX) and facility-of-origin (Zion vs. Parnassus). (**E**) CAS of Zion (N=3) and Parnassus (N=6) female SKG mice treated with MTX three times weekly in the absence of co-housing. (**F**) Comparison of CAS in two experiments of mice treated with MTX over a 14-day time frame. (**G**) Comparison of CAS AUC in two experiments. Results of Student’s t-test are reported. Error bars are SEM.

Because we observed variation in treatment response in this pilot experiment, we stratified the data by facility-of-origin (**Figure 6C**). Remarkably, we saw that while arthritis development was similar for both groups of mice in the VEH arm (**Figure 6C, left panel**), the response to MTX was significantly better in Zion-reared mice compared to Parnassus-reared mice (**Figure 6C, right panel**); evaluation of the AUC of the arthritis curves also supported this finding (**Figure 6D**). This difference was reproduced in a second set of mice that were not co-housed (“Experiment 2”). Mice reared at Zion showed significantly lower arthritis scores on Days 7 through 14 (**Figure 6E, left panel**), which is also reflected in the arthritis AUC (**Figure 6E, right panel**). Together, these results suggest that facility-of-origin of the mice impacted treatment response to MTX. Of note, this murine phenotype recapitulates what is seen in RA: patients possess similar arthritis severity but can vary significantly in their response to MTX. These findings demonstrate that the SKG arthritis model responds to oral MTX and that it can be used to dissect mechanisms underlying variable treatment response.

We next tested for differences between the gut microbiomes of mice reared in these two facilities. We studied two sets of mice to understand the impact of co-housing: “representative” mice that did not undergo co-housing (“Representative”) as well as mice that were co-housed for 2 weeks prior to arthritis induction and treatment (Experiment 1). We performed 16S rRNA gene amplicon sequencing (16S-seq) on stool samples from Zion-reared SKG mice (N=5), Parnassus-reared SKG mice (N=4), and mice from our pilot co-housing experiment prior to the start of any treatments (Experiment 1, Zion-reared N=7; Parnassus-reared N=9). Alpha diversity was significantly higher in Zion-reared mice (**Supplementary Figure 5A, left panel**) compared to Parnassus-reared mice. This difference was effectively eliminated by co-housing (**Supplementary Figure 5A, right panel**), with Parnassus-co-housed mice acquiring multiple amplicon sequence variants (ASVs, representative of individual species, **Supplementary Table 2**). Community composition (beta diversity) differed by facility-of-origin in the “Representative” mice (ANOSIM p=0.013, PERMANOVA p=0.017; **Supplementary Figure 5B**, open circles). In the co-housed mice, this difference persisted, albeit in a reduced form (ANOSIM p=0.01, PERMANOVA p=0.054; **Supplementary Figure 5B**, closed circles). We next identified differentially abundant taxa. There were no significant phylum-level differences by facility-of-origin in either group. While 123 ASVs were differentially abundant in the “Representative” mice (with 109 higher in Zion and 14 higher in Parnassus), after 2 weeks of co-housing, only 8 ASVs were differentially abundant by facility-of-origin (with 6 higher in Zion and 2 higher in Parnassus, **Supplementary Figure 5C**). These findings confirm that the microbiome of Zion- and Parnassus-reared mice differed and that that co-housing facilitated transfer of microbes from Zion mice to Parnassus mice. Given that we observed that treatment response to MTX was better in Parnassus-cohoused-with-Zion mice compared to Parnassus-reared mice (**Figure 6F-G**), this suggests that transfer of microbes from Zion to Parnassus improved treatment response in this mouse model.

### MTX response is linked to MTX metabolism *in vivo*

Next, we tested whether the mechanism for differential MTX response was linked to microbial MTX metabolism and modulation of MTX PK *in vivo*. We re-derived the SKG model in our gnotobiotic facility. This enables us to selectively examine the contribution of gut microbes and control for other factors. We colonized GF SKG mice with cecal microbiota from either Parnassus-vs. Zion-reared mice (N=5-6 per colonization group). We found that peak plasma concentrations of MTX were higher in mice colonized with Parnassus microbiota (**Figure 7A-B**). The ratio of 7-OH-MTX to MTX (7-OH-MTX_AUC_/MTX_AUC_) was numerically higher in Parnassus-colonized mice (**Figure 7D**). Microbial metabolism was also dependent on colonization state: DAMPA levels and AUC were higher in mice colonized with Parnassus microbiota (**Figure 7A-C**). pMABG levels were numerically higher in Parnassus-colonized mice, but we were unable to detect a significant difference in the plasma (**Figure 7A-C**).

**Figure 7.**
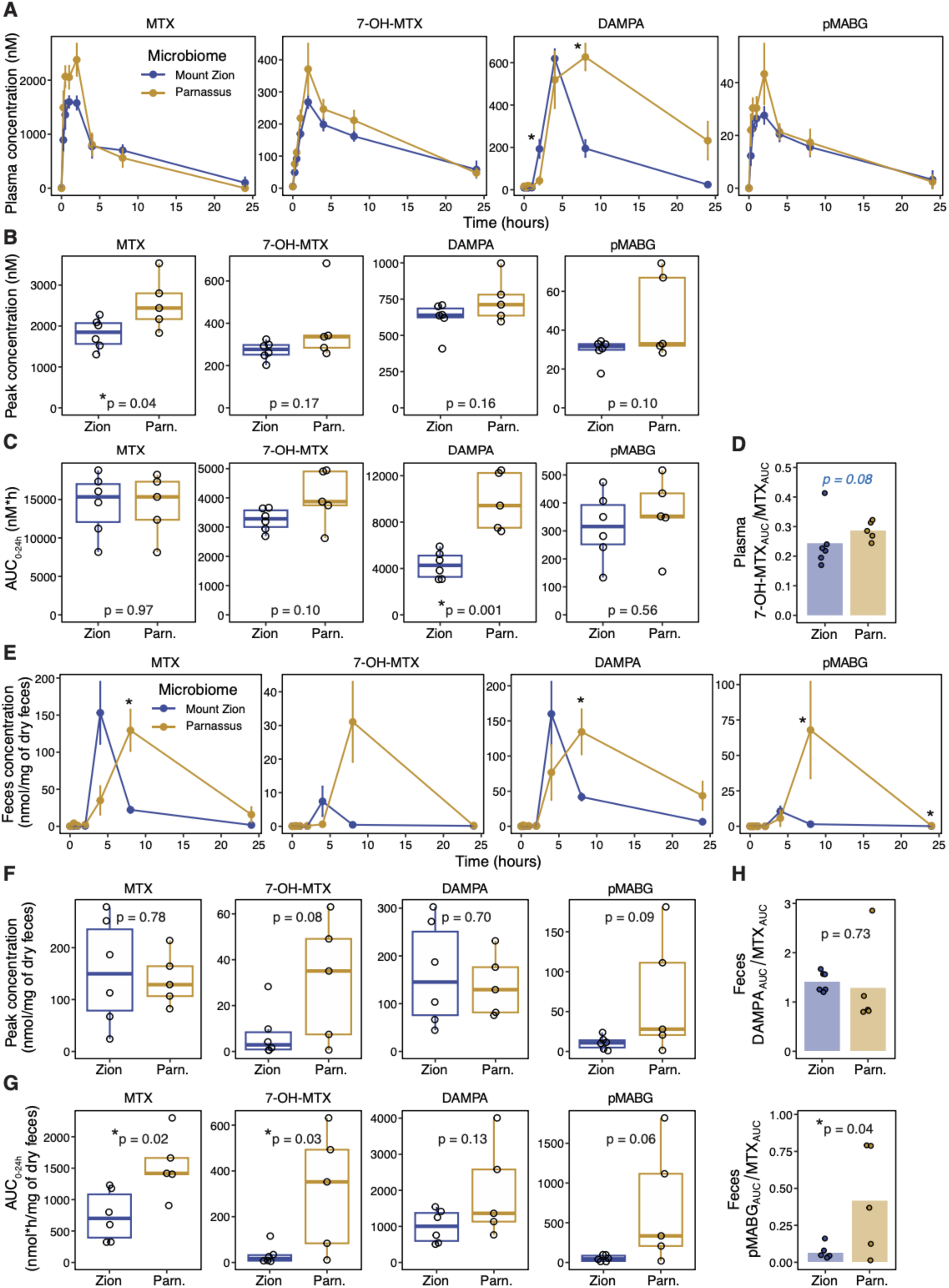
MTX response in a model of inflammatory arthritis is linked to microbial metabolism of MTX. (**A**) Plasma concentration of MTX and its metabolites were quantified by LC-MS in female GF SKG mice colonized with either Zion or Parnassus (Parn.) microbiota (N=6 per colonization state). After a one-time oral administration of MTX 50 mg/kg, multiple plasma samples were taken from each mouse over a 24 h time course. (**B**) Peak concentration and (**C**) AUC were compared between Zion and Parnassus colonized mice. (**D**) Ratio of AUC of 7-OH-MTX to MTX in the plasma over 24 h. (**E**) Stool concentrations of MTX and its metabolites were quantified at multiple time points, and (**F**) peak stool concentrations, (**G**) AUC and (**H**) product-substrate ratios of DAMPA (top panel) and pMABG (bottom panel) relative to MTX are shown. Results from Student’s t-test (regular black) or Wilcoxon test (italic blue, data are non-normal by Shapiro-Wilk test) are reported. Error bars are SEM.

In the stool, MTX levels and AUC were significantly higher in Parnassus-colonized mice (**Figure 7E-G**) suggesting greater intestinal elimination. AUC of 7-OH-MTX was significantly higher in Parnassus-colonized mice (**Figure 7E-G**). Similarly, fecal DAMPA and pMABG levels were significantly higher in Parnassus-colonized mice (**Figure 7E**) as was the product-substrate ratio of fecal pMABG to MTX, suggesting increased microbial metabolism.

Taken together, this suggests that while they may permit increased MTX absorption relative to Zion-microbiota, Parnassus-microbiota led to increased microbiota- and host-dependent MTX metabolism as detected in the blood and stool. These findings suggest that microbiota-dependent metabolism of MTX contributes to MTX PK and is linked to clinical outcomes in inflammatory arthritis.

## Discussion

Large-scale *in vitro* screens have identified multiple therapeutic drugs that are metabolized by human gut microbiota^3,4^, but the pharmacologic and clinical repercussions of these microbial transformations remain largely unknown. Here, we advance our understanding of how the gut microbiome interacts with a key anti-inflammatory drug, MTX, a WHO essential drug used by multiple wide-ranging clinical disciplines. MTX has been extensively studied for over 50 years. By taking a microbiologic lens, we uncover novel pharmacology that fundamentally transforms our understanding of how the gut microbiome interacts with MTX to shape therapeutic outcomes. We show that the gut microbiome contributes to MTX PK *in vivo* through both direct and indirect mechanisms, that natural variation in human microbiota leads to differential MTX PK profiles, that multiple human-associated microbes metabolize MTX into known (DAMPA) and novel (pMABG and 6-MPDA) metabolites, and that microbial metabolism *in vivo* in mice is linked to treatment outcomes. While it has long been known that there is extensive interindividual variation in the bioavailability of MTX^31^, the mechanisms underpinning this variation remain incompletely elucidated. Here, we provide evidence from multiple orthogonal systems (*in vitro*, *ex vivo*, and *in vivo* mouse studies) to suggest that the gut microbiome plays a pivotal role. Given that the microbiome is modifiable through diet, probiotics, transplant, and other strategies, these findings provide support for targeting the microbiome to maximize the efficacy of this important anti-inflammatory drug.

Our findings place the microbiome on the map as a determinant of MTX response. While one of the very first MTX metabolites to be discovered was microbiota-produced DAMPA, the extent of microbial metabolism was reported to be very minimal^24^; our findings run counter to this. This may be because these early studies were performed in cancer patients receiving intravenous MTX, and DAMPA was quantified in urine, not the gut, where it is produced^22^. Given that MTX is administered orally for autoimmunity, these early findings in cancer patients may not generalize. Our results suggest that MTX undergoes significant metabolism by the gut microbiota, into both DAMPA and pMABG, and that the highest levels of these are found in the gut, where they are produced. Importantly, we show that DAMPA and pMABG are found in the blood as well. In some of our studies (mice colonized with their native microbiota), we find that levels of DAMPA approximate concentrations achieved by host-produced metabolites, namely 7-OH-MTX, suggesting that DAMPA may be as important as its host-produced counterpart and should not be considered a minor metabolite. Further, the microbiota regulates liver metabolism of MTX into 7-OH-MTX, thereby indirectly impacting MTX pharmacology. Indeed, prior studies show that AO, the enzyme responsible for 7-OH-MTX production, is upregulated in colonized mice relative to GF mice^40^ and that interindividual variation in AO in humans impacts 7-OH-MTX production^28^. Given the extensive variation in gut microbiotas across patients, we suspect that the gut microbiome plays a critical role in MTX bioavailability and drug metabolism. Developing microbiota-directed interventions will allow patients to derive maximal benefit from this central drug. Future studies are needed to understand natural variation in the production of these metabolites and their association with response in patients, the physiologic impacts of microbiota-produced metabolites on the host and its commensal microbiome, as well as how best to specifically modulate the human microbiome to improve drug response.

Our findings showcase how taking a microbiome perspective leads to unexpected pharmacological insights. For instance, we were surprised to discover novel MTX metabolites given that the pharmacology of MTX had been investigated for decades. Stable isotope labeling combined with *in vitro* microbiologic assessment allowed us to identify novel MTX metabolites; GF mouse studies revealed that one of these novel metabolites, pMABG, is exclusively microbially produced. For the smaller metabolite, 6-MPDA, though we were able to detect this in the *in vitro* stable isotope study, we had difficulty quantifying it in our mouse experiments, in either the blood or the feces. This may be due to its small molecular weight, suggesting that we need to further optimize methods to quantify these small molecules. Another possibility is that it is produced and immediately shuttled into host or microbial biosynthetic pathways. As for pMABG, no living organisms have been reported to produce this compound. Interestingly, the folate analog, p-aminobenzoylglutamatic acid (pABG), is produced and even used by organisms as a nutrient source^41^. Thus, pathways that produce folate-derived pABG may also act on MTX to produce pMABG. The identity of these enzymes remain unknown; indeed, there are still many unknowns with respect to folate metabolism^42^. Similar to the way in which MTX opened up a “Golden Era” in the discovery of folate metabolism^43^, microbial metabolites of MTX may uncover novel pathways in microbes that are of fundamental importance. While the identity of these microbial enzymes remains to be determined, our findings reveal that human gut microbes produce novel MTX metabolites *in vitro* and *in vivo*, and that these compounds, which are produced in the gut, can enter into circulation.

Finally, our work showcases the translational implications of basic science studies focused on *in vitro* microbial drug metabolism^3,4^ and suggests that microbiome modification can be plausibly leveraged to improve drug efficacy *in vivo*. Taken together with our previous findings that microbiome function and *ex vivo* metabolism is associated with clinical response to MTX^19^, the findings reported here in preclinical models provide strong evidence for the role of the microbiome in shaping treatment outcomes for patients taking MTX.

## Limitations

We found that SPF mice reared in two different campus facilities experienced different responses to MTX, that response was transferrable by co-housing, and that GF colonized with these different microbiotas showed different PK profiles as well as metabolite production. A natural extension would be to evaluate MTX response in an arthritis experiment with GF colonized mice with Zion-vs. Parnassus-microbiota. However, when we performed power calculations in preparation for this experiment, we learned that it would be technically challenging and costly because arthritis scores are lower in gnotobiotic mice for multiple reasons, leading to “floor effects”^44,45^. For instance, it is well-known that gnotobiotic SKG mice experience significantly reduced arthritis in the gnotobiotic setting^35,38^. Moreover, in the setting of MTX treatment, both colonization arms (i.e., Zion vs. Parnassus) experience further reductions in arthritis scores; detecting a difference between colonization states will require large numbers of gnotobiotic mice (upwards of 30-40 per colonization group) and expensive isolators. While these logistical and financial challenges may be overcome in future work requiring extensive optimization, the preponderance of evidence presented in this and previous reports^19,20^ serve to support the main conclusions presented here. Another limitation is that while we were able to detect pMABG in the blood and stool of mice and in *ex vivo* cultures of RA patients incubated with MTX, the extent to which this metabolite is found in human circulation is unknown.

## Conclusion

In conclusion, we have shown that the human gut microbiome metabolizes a key anti-inflammatory drug, MTX, and shapes the pharmacology of this drug *in vivo*. The microbiome directly metabolizes MTX and also affects host metabolic pathways that shape drug pharmacology. We identified novel metabolites of MTX, one of which likely is a major metabolite found in high abundance in the stool. This metabolism is associated with treatment response in a mouse model of inflammatory disease. Taken together with prior human studies^19,20^, these findings provide a strong foundation for future microbiome-directed interventions to improve MTX outcomes. Further, we provide a framework to investigate additional drugs that undergo microbial metabolism and evaluate impacts *in vivo*. These findings support the notion that microbial enzymes that are beneficial to the host for nutrient harvest (e.g., folate metabolism) may also lead to unintended metabolism of therapeutic drugs and suggest that the metabolic capacity of the microbiome should be considered when personalizing therapies for patients.

## Material and methods

### RESOURCE AVAILABILITY LEAD CONTACT

Further information and requests for resources and reagents should be directed to and will be fulfilled by the Lead Contact, Renuka R. Nayak

### MATERIALS AVAILABILITY

We rederived the SKG mouse model in the UCSF Gnotobiotic Core facility during the course of this study. Multiple human gut bacterial strains were isolated and are available upon request.

### DATA AND CODE AVAILABILITY

Upon publication, all sequencing data generated in the preparation of this manuscript will be deposited in NCBI’s Sequence Read Archive (SRA) under a to-be-determined Bioproject ID.

### EXPERIMENTAL MODEL AND SUBJECT DETAILS

#### Mouse experiments

##### Animal strains and care

GF and SPF C57BL/6J and BALB/c SKG mice (males and/or females, ages 6-13 weeks) were born at UCSF. SPF mice were kept in SPF conditions in the Animal Barrier Facility at UCSF, while GF mice were maintained within the UCSF Gnotobiotics Core Facility (gnotobiotics.ucsf.edu) and housed in gnotobiotic isolators for the duration of the experiment (Class Biologically Clean). The mice were housed at temperatures ranging from 19 °C to 26 °C and humidity ranging from 30 to 70% with a 12 h/12 h light/dark cycle. Mice had *ad libitum* access to water and standard chow diet (Lab Diet, cat # 5058, USA) in the Parnassus SPF facility and (Lab Diet, cat # 5053, USA) in the Mount Zion (MTZ) SPF facility, while standard autoclaved chow diet (Lab Diet, cat # 5021, USA) was used in the gnotobiotic facility.

Similarly, GF and SPF SKG mice were born at UCSF. GF SKG mice were rederived by Cesarean delivery into the University of California, San Francisco (UCSF) Gnotobiotic Core Facility and maintained in GF isolators for breeding. Stool pellets from isolators in the Gnotobiotic Core Facility were screened monthly by PCR for 16S bacterial and 18S (Internal transcribed spacer) fungal ribosomal sequences. Mice were randomly housed based on sex.

Several independent experiments were carried out and, for each experiment, mice were randomly assigned to groups. Key details for each mouse experiment can be found in **Supplementary Table 3**. All mouse experiments were approved by and conformed to the standards of the University of California San Francisco Institutional Animal Care and Use Committee.

##### Mice colonization

For humanization of GF mice with a human microbiome, mice were colonized with stool from treatment-naïve human donors with RA that were collected as part of a prior study^19^. Human stool was diluted 1:10 g/mL in prereduced sterile PBS + 0.05% cysteine and homogenized in an anaerobic chamber (Coy Laboratory Products) using pre-equilibrated reagents and supplies. Insoluble material was separated from supernatant by centrifugation at 50 *g* for 1 min. Aliquots of supernatant (100 μL per mouse) were gavaged into mice. Mice were colonized for at least 2 weeks before initiation of treatment with MTX.

For colonization of mice with murine microbiota, cecal contents from SPF BALB/c SKG mice (either Zion or Parnassus) were harvested after euthanasia. The gastrointestinal tract was excised and regions proximal and distal to the cecum were tied off with dental floss. This was brought into an anaerobic chamber where the cecum was opened and the contents were placed into a conical. Cecal contents were diluted 1:10 g/mL in prereduced sterile PBS + 0.05% cysteine and homogenized in an anaerobic chamber using pre-equilibrated reagents and supplies. Insoluble material was separated from supernatant by centrifugation at 50 *g* for 1 min. Aliquots of supernatant (100 μL per mouse) were gavaged into GF BALB/c SKG mice. Mice were colonized for at least 2 weeks before initiation of treatment with MTX.

#### Human subjects

Human subject samples were used for colonizing GF mice with human microbiota and also for *ex vivo* profiling. These patients were enrolled either as part of a prior study^19^ and as part of an ongoing study NYU-UCSF collaboration. Consecutive patients from the New York University Langone Medical Center’s rheumatology clinics and offices were screened for the presence of RA based on ACR criteria^46^. After informed consent was signed, each patient’s medical history (according to chart review and interview/questionnaire), diet, and medications were determined. A screening musculoskeletal examination and laboratory assessments were also performed or reviewed. All RA patients who met the study criteria were offered enrollment. The criteria for inclusion in the study required that patients meet the American College of Rheumatology/European League Against Rheumatism 2010 classification criteria for RA^46^, including seropositivity for rheumatoid factor and/or anti–citrullinated protein antibodies, and that all subjects be age 18 years or older. New-onset RA was defined as disease duration of a minimum of 6 weeks and up to 6 months since diagnosis, and absence of any treatment with disease-modifying anti-rheumatic drugs, biologic therapy or steroids (ever). The exclusion criteria applied to all groups were as follows: recent (<3 months prior) use of any antibiotic therapy, current extreme diet (e.g., parenteral nutrition or macrobiotic diet), known inflammatory bowel disease, known history of malignancy, current consumption of probiotics, any gastrointestinal tract surgery leaving permanent residua (e.g., gastrectomy, bariatric surgery, colectomy), or significant liver, renal, or peptic ulcer disease. This study was approved by the Institutional Review Board of New York University School of Medicine protocols. All new onset rheumatoid arthritis patients were recruited using established protocols from a previously described study^19^. Pre-treatment stool samples were collected at baseline. Clinical and demographic data was de-identified and recorded in RedCap by the designated study personnel.

### METHOD DETAILS

#### Methotrexate administration and pharmacokinetic studies

For each experiment evaluating the PK profile of MTX, a single dose of 50 mg/kg (Pharmaceutical-grade, Fresenius Kabi, NDC 63323-122-50, cat # 102250) was administered by oral gavage (PO), except for the route of administration experiment where half of the mice were treated by intraperitoneal injection (IP). MTX was dissolved at 50 mg/mL in a vehicle (VEH) containing 66% of PBS 1X and 33% of NaOH 0.1 M. MTX solution was filtered through a 0.22 μm filter. Blood samples were collected from the tail of awake freely moving mice at the following time points: 0 (before treatment), 15, 30, 60, 120, 240, and 440 min after treatment; for blood collection, we used microhematocrit capillary tubes (Fisher #22-362-566). For some experiments, a 24-h time point was also included. After centrifugation at 3,500 *g* for 15 min at 4 °C, plasma was collected. In parallel to blood sampling, stool samples were collected whenever possible. Plasma and stool samples were stored at −80 °C for targeted metabolomics (*see below*). From the concentration–time curve obtained from each individual mouse, *max* and *auc* functions from the ‘MESS’ package (v0.5.12) in R were used to calculate individual peak concentration and AUC values. Of note, a value of 0 nmol/mg of dry feces was inferred for a metabolite when no stool sample could be obtained from a mouse for the initial 0 min time point and all other mice showed a value of 0.

#### Targeted metabolomics for PK studies

##### Plasma

Plasma quantification of MTX and its metabolites was performed by targeted metabolomic analysis using liquid chromatography-tandem mass spectrometry (LC–MS). Samples were analyzed using a SCIEX ExionLC autosampler (Shimadzu SIL-30AC series) in sequence with a hybrid triple quadrupole – linear ion trap mass spectrometer (either SCIEX 6500 QTRAP or a SCIEX 7500 QTRAP, depending on machine availability) (Shimadzu, USA). 5 μL of each sample was diluted with 45 μL HPLC-grade H₂O and extracted with 200 μL of HPLC-grade methanol (containing either aminopterin or 2-amino-3-bromo-5-methylbenzoic acid as internal standard). Samples were incubated for at least 30 min at −20 °C before being centrifuged at 13,000 rpm for 10 min. 100 μL of supernatant was dried using a SpeedVac concentrator (Savant, cat # DNA 120) and then reconstituted in 100 μL of HPLC-grade H₂O. To minimize fluctuations in pressure and clogging, a filtering step was implemented using a 0.2 µm PVDF hydrophilic membrane filter plate (Corning, cat# 3505) at 3,200 rpm for 5 min.

##### Feces

Feces quantification of MTX and its metabolites was performed similarly to plasma samples (using the SCIEX 7500 QTRAP). Fecal samples were lyophilized; wet and dry weight was recorded before and after lyophilization. Dried feces were extracted with 500 uL of extraction solvent containing 80% methanol in H₂O and 0.1% formic acid (with 2-amino-3-bromo-5-methylbenzoic acid as internal standard). Samples were homogenized using the Precellys 24-bead homogenizer pre-cooled with dry ice at 6,400 rpm for 3 cycles of 20 s. Samples were centrifuged at 4 °C for 15 min at 16,000 rpm. 400 uL of supernatant was transferred into new tubes, and speed vacuumed until completely dry. Dried samples were re-suspended in 200 uL of HPLC-grade H₂O and a 1:10 dilution was performed. To minimize fluctuations in pressure and clogging, a filtering step was implemented using a 0.2 μm PVDF hydrophilic membrane filter plate (Corning, cat# 3505) at 3,200 rpm for 5 min.

##### MRM method development

Multiple reaction monitoring (MRM) was established for each compound using concentrated pure standards directly on a SCIEX 7500 or 6500 Triple Quad Mass Spectrometer using a syringe pump model. Intense peaks were used to identify the molecular fragmentation (Q3 mass), and collision energy (CE) ramping was used to determine the optimal detection of the standards on the instrument, with reported optimization values referenced in **Supplementary Table 4**.

Two different methods were used to quantify MTX and its metabolites. Method A was used initially to quantify MTX, 7-OH-MTX and DAMPA. Method B was developed in order to quantify these as well as the novel MTX metabolites: pMABG and 6-MPDA. 5 μL of reconstituted samples were injected into the system. Analytes were chromatographically separated using either a Synergi 4 µm Fusion-RP 80 Å, 2 x 50 mm HPLC column (Phenomenex, cat# 00B-4424-B0) for method A, or a Kinetex F5 2.6 µm, 2.1 × 150 mm LC column (Phenomenex, cat # 00F-4723-AN) for method B.

In method A, a mobile phase scheme consisting of [A] H_2_O + 0.1% formic acid and [B] methanol + 0.1% formic acid was used. LC flow was set to a constant rate of 600 μL/min, and the timed linear gradient program consisted of: time = 0 min, 5% B, time = 1 min, 5% B, time = 5 min, 40% B, time = 5.25 min, 100% B, time = 5.75 min, 100% B, time = 6 min, 5% B, and time = 8 min, 5% B. Data was collected in positive polarity mode using MRM with the following source parameters: curtain gas (CUR) = 35 psi; nebulizer gas (GS1) = 50 psi; heater gas (GS2) = 70 psi; collision gas = 9; source temperature (TEM) = 450 °C; ion spray voltage (ISV) = 1,500 V. Other method parameters used include a cycle time of 0.525 s, dwell time of 100 ms/MRM pair, settling time = 0 ms, and pause time of 5.007 ms.

In method B, a mobile phase scheme consisting of [A] H_2_O + 0.1% formic acid and [B] acetonitrile + 0.1% formic acid was used. LC flow was set to a constant rate of 200 μL/min, and the timed linear gradient program consisted of: time = 0 min, 0.2% B, time = 1 min, 0.2% B, time = 2 min, 10% B, time = 4 min, 25% B, time = 6 min, 98% B, time = 8 min, 98% B, time = 8.1 min, 0.2% B, and time = 10 min, 0.2% B. Data was collected in positive and negative polarity modes using MRM with the following source parameters: CUR = 35 psi; GS1 = 50 psi; GS2 = 70 psi; collision gas = 9; Temperature = 450 °C; ISV = 2,200 V (positive) and −2,200 V (negative). Other method parameters used include a cycle time of 0.435 s, dwell time of 10 ms/MRM pair, settling time = 15 ms, and pause time of 5.007 ms.

Raw data was processed using a built-in SCIEX OS software package (version 2.1.6.59781) for peak picking, alignment, and quantitation. Peak areas of the standard curves were converted to concentrations at ng/mL using linear regression, then expressed as nM (plasma samples) or nmol/mg of dry feces (stool samples).

#### Stable isotope profiling

Labeled methotrexate was purchase (d3-MTX, Santa Cruz # sc-218707) along with unlabeled MTX. Stock concentrations were dissolved in DMSO at a concentration of 90 mg/mL. Labeled MTX, unlabeled MTX, or DMSO was further diluted to achieve a final concentration of 100 ug/mL into the experimental plate. For each of *Bacteroides thetaiotaomicron* DSM 2079, *Clostridium asparagiforme* DSM 15981, and *Clostridium symbiosum* DSM 934, a single colony of each isolate was subcultured in Bacto Brain Heart Infusion media (BD Biosciences, 37 g/L) supplemented with L-cysteine-HCl (0.05%, w/v), menadione (1 µg/mL), and hemin (5 µg/mL) (referred to hereafter as BHI+) for 48 h in an anaerobic chamber at 37 °C with an atmosphere composed of 2-3% H2, 20% CO2, and the balance N2. This subculture was diluted down to an OD600 of ∼0.1, which was then further diluted 100-fold, and then used to inoculate the experimental plate containing the drug. Sterile controls were also tested for each drug. After 72 h, 100 uL aliquots were centrifuged at 1,900 rcf for 5 min at room temperature, and 40 uL of the supernatant was submitted for global LC-MS profiling. Each drug-bacterial species pair and sterile control was tested in quadruplicate.

#### Global profiling of isotope labeled *in vitro* monocultures

Samples were thawed and combined with 80 µL of LC-MS-grade methanol: acetonitrile (1:1 v/v and 1µM chlorpropamide) with approximately 10 µL of 0.1 mm zirconia beads. Samples were then thoroughly homogenized in BeadBlaster 24 Microtube Homogenizer 2 x 30 s at 4,260 rpm. Samples were incubated at −20 °C for 1 h. Then samples were vortexed and centrifuged at 12,000 g at 4 °C for 15 min. Supernatant was transferred to autosampler vials for LC-MS analysis.

Samples (5uL) were separated by reverse phase HPLC using a Prominence 20 UFLCXR system (Shimadzu, Columbia, MD) with a Waters (Milford, MA) BEH C18 column (100 mm x 2.1 mm 1.7 um particle size) maintained at 55 °C and using a 20 min aqueous acetonitrile gradient, at a flow rate of 250 uL/min. Solvent A was HPLC-grade H₂O with 0.1% formic acid and Solvent B was HPLC-grade acetonitrile with 0.1% formic acid. The initial conditions were 97% A and 3% B, increasing to 45% B at 10 min, 75% B at 12 min where it was held at 75% B until 17.5 min before returning to the initial conditions. The eluate was delivered into a 5600 (QTOF) TripleTOF using a Duospray™ ion source (all Sciex, Framingham, MA).The capillary voltage was set at 5.5 kV in positive ion mode with a declustering potential of 80 V. The mass spectrometer was operated with a 100 ms TOF scan from 50 to 1000 *m/z*, and 16 MS/MS product ion scans (100 ms) per duty cycle using a collision energy of 50 V with a 20 V spread.

#### RA patient fecal sample *ex vivo* profiling

All work was carried out in an anaerobic chamber. For each patient, stool was aliquoted into a pre-equilibrated cryovial, diluted in pre-reduced PBS at 10 mL/g of stool, and vortexed to homogenize the sample. The sample was spun at ∼20 *g* for 1 min on a mini-centrifuge to facilitate settling of sediments, and the sediment-free supernatant (“fecal slurry”) was then aliquoted into a new pre-equilibrated cryovial for evaluation of *ex vivo* growth. Growth was evaluated by inoculating liquid BHI+ with 1:50 dilution of this fecal slurry. Samples were treated with MTX 100 μg/mL or the vehicle control and incubated with shaking for 24 h at 37 °C, and then stored at −80 °C for future profiling. For LC-MS profiling, samples were thawed, centrifuged at 4 °C at 13,000 g for 5 min. 200 μL of supernatant was mixed with 800 μL of LC-MS-grade methanol: acetonitrile (1:1 v/v) and incubated at −20 °C for 1 h. Samples were vortexed, and then centrifuged at 15,000 g for 15 min at 4 °C. Supernatant was transferred to autosampler vials for targeted metabolomics.

#### *Ex vivo* fecal sample targeted metabolomics

The analysis was performed using a Nexera 20 HPLC system (Shimadzu) coupled to a TripleTOF 5600 mass spectrometer (Sciex) equipped with a DuoSpray ion source, operating in electrospray ionization (ESI) mode with positive ion detection. Chromatographic separation was achieved on a Waters XSelect HSS T3 column (2.1 × 100 mm, 2.5 µm particle size) maintained at 40.0 °C. The mobile phase was delivered at a flow rate of 0.200 mL/min. Solvent A consisted of LC/MS-grade water with 0.1% formic acid, and Solvent B was LC/MS-grade acetonitrile containing 0.1% formic acid. The gradient elution program was as follows: 0.00-5.00 min, 5.0% B; 9.00 min, 90.0% B; 14.00 min, 100.0% B; 14.50–20.00 min, 5.0% B. The injection volume was 2.0 µL. Mass spectrometric data (MS1 and MS2) were acquired using a declustering potential (DP) of 70 V. MS1 data were collected over a mass range of 100–1000 Da. The ion source parameters were set as follows: ion spray voltage (IS), 5500 V; curtain gas (CUR), 30 psi; nebulizer gas (GS1), 60 psi; heater gas (GS2), 50 psi; and source temperature (TEM), 500 °C. High-resolution multiple reaction monitoring (MRM) was employed with a total cycle time of 500 ms and unit resolution in Q1. The following precursor ions and their respective collision energies (CE) were used for MS2 analysis:

- pMABG (m/z 281.13, CE 55 eV, 8.50 min)
- MTX (m/z 455.20, CE 50 eV, RT 8.35 min)
- DAMPA (m/z 326.15, CE 34 eV, RT 8.60 min)

Data were analyzed using MultiQuant software (Sciex, v3.0.3) with the MQ4 quantitation algorithm. Quantification was performed against a standard curve of authentic standards at nine concentrations ranging from 1 nM to 1 μM.

The following fragment ions were used for quantitation:

- MTX: m/z 308.13
- DAMPA: m/z 176.08
- pMABG: m/z 134.06

Masses were integrated with a 10 mDa tolerance.

#### Isolation of microbes from patient stool samples

Bacterial isolates were harvested from MTX-naïve RA patient fecal slurries prepared as previously described^19^.For isolate harvesting, fecal slurries were diluted 1:20 in BHI+ media then serially diluted 1:10 in BHI+ media 7 times. Serial dilutions were streaked on BHI+ agar plates. To enrich for *Firmicutes*, another set of serial dilutions were exposed to aerobic conditions and 70% (v/v) ethanol for 4 h prior to streaking on BHI+ agar plates supplemented with taurocholate as previously described^34^. Isolates were grown at 37 °C for 48 h in an anaerobic chamber. Unique colony morphologies were identified and picked for growth in liquid BHI+ media at 37° C for 48 h before reconstitution in 2:1 of 80% (v/v) glycerol and placement in −80 °C for long-term storage.

#### Bacterial DNA extraction and full-length 16S rRNA sequencing

Full-length Sanger sequencing of the 16S rRNA gene was performed by either extracting DNA from turbid, 48-h cultures; verification was done by colony PCR. *<u>Liquid cultures</u>*: For each isolate, bacterial liquid cultures (2 mL) incubated for 48 h in an anaerobic chamber were pelleted by centrifugation for 5 min at 2,000 rcf and the supernatant was discarded. The bacterial pellet was resuspended in 200 uL Instagene Matrix (Cat # 732-6030) and incubated at 56 °C and 100 °C for 30 min and 8 min respectively. Samples were vortexed before and after each incubation. *<u>Colony</u> <u>PCR</u>*: After streaking glycerol stocks and incubating for 48 h at 37 °C under anaerobic conditions, an individual colony was picked and placed in a PCR strip-tube containing PCR master mix. The PCR master mix consisted of 25 uL Amplitaq gold (Life Tech/Thermo Fisher 438881), 1 uL forward primer (8F, 5’-AGAGTTTGATCCTGGCTCAG-3’), 1 uL reverse primer (1542R, 5′-AAGGAGGTGATCCAGCCGCA-3’), and 18 uL molecular grade H₂O per reaction. PCR amplification was as follows: 10 min at 95 °C, 29x (30 s at 95 °C, 30 s at 50 °C, 30 s at 72 °C), and ending at 4 °C.

PCR purification was achieved by using a PureLink PCR Purification Kit (Invitrogen K3100-01) or Axygen AxyPrep Mag kit (#MAG-PCR-CL-5) according to the manufacturer’s instructions. Purified DNA was quantified using Nanodrop (Thermo Scientific NanoDrop 2000 Spectrophotometer). In PCR strip-tubes, 10 uL of the mix template and 5 uL of each primer (either F or R) was mixed and submitted to GENEWIZ for Sanger Sequencing. Forward and reverse reads were combined into a contig that was analyzed using BLAST (<u>BLAST: Basic Local Alignment Search Tool (nih.gov)</u>) where the highest “percent identity” was chosen for species identification.

#### MTX metabolism screen

RA-associated strains were grown for 48 h in an anaerobic chamber at 37 °C and were subcultured at a 1:50 dilution (2 uL of bacterial culture in 98 uL culture media) in a 96-well U bottom plate. Bacteria were grown in BHI+ with MTX 50 ug/mL or vehicle control (DMSO). After 72 h of incubation at 37 °C, the plates were placed in the −80 °C freezer until extraction.

#### HPLC-based quantification of MTX levels of *in vitro* studies

For bacterial samples, plates were spun down at 2,400 rpm 10 min to pellet cells. 30 uL of the bacterial supernatant was extracted with 150 uL of HPLC-grade methanol. The samples were incubated at −20 °C for 30 min. After incubation, the plates were centrifuged at 3,000 rpm for 30 min. After centrifugation, 60 uL of supernatant from each well was transferred into 200 uL plastic inserts in HPLC glass vials. HPLC-grade methanol was used as blank.

The samples were profiled by reversed phase HPLC using an Agilent 1260 Infinity II system with a Kinetex 2.6 um C18 column and a 8 min gradient, at a flow rate of 0.6 mL/min. For the mobile phase, solvent A was 0.1% formic acid and solvent B was 100% HPLC-grade methanol. The initial condition was 90% B, decreasing to 70% B by 1 min, 0% B by 7 min, and 90% B by 7.5 min where it was held until 8 min. Authentic standards of MTX were used for peak identification and a standard curve was generated for quantification (100, 50, 25, 12.5, 6.25 ug/mL). When MTX depletion is reported (usually as a percentage), we normalize each sample to sterile controls on the same plate. Occasionally, we noted MTX percentages > 100%, which we hypothesized could be due to evaporation effects.

#### Validation of MTX metabolism for bacterial strains *in vitro*

Glycerol stocks from harvested RA-associated strains were streaked on BHI+ agar plated in an anaerobic chamber. A single colony was subcultured in BHI+ liquid media for 48 h at 37 °C. This subculture was diluted down to an OD600 of 0.08 - 0.1, which was then further diluted 100-fold, and then exposed to MTX at a final concentration of either 50 ug/ml in quadruplicate along with vehicle controls (DMSO). Plates were incubated at 37 °C for 72 h. An Eon Microplate Spectrophotometer (BioTek Instruments, Inc) was used to measure OD600 at 0 and 72 h to confirm bacterial growth. Data was exported using the Gen5 (Version 3.12) software. After 72 h, aliquots were prepared for extraction and profiling on HPLC (*see above,* “HPLC-based quantification of MTX levels of *in vitro* studies”) and LC-MS platforms.

#### LC-MS analysis of *in vitro* isolates

A single sample of each bacterial species incubated with MTX (50 - 100 ug/ml) or sterile control was selected for profiling on LC-MS. From each sample, 60 uL of supernatant was transferred and stored at −20 °C. Based on MTX depletion and detection of novel peaks on HPLC, a sample from each bacterial species was selected: 60 uL was dried and reconstituted in 1 mL HPLC-grade H_2_O. The samples were filtered using a 0.2 μm PVDF hydrophilic membrane filter plate (Corning, cat# 3505) at 3,200 rpm for 5 min. 100 uL of the samples were transferred into glass inserts and vials for LC-MS profiling. LC-MS targeted metabolomics (*see above,* “Targeted metabolomics for PK studies” method B) was used for quantifying MTX and its metabolites with a standard curve of authentic standards at concentrations ranging from 0.2 to 200 ng/mL.

#### SKG arthritis experiments

To evaluate the therapeutic effect of MTX on an arthritis mouse model, BALB/c SKG mice were given once 100 uL of zymosan emulsion (10 mg/mL) through IP injection to induce arthritis. Zymosan (Sigma, #Z4250-1G) was emulsified in sterile 1X PBS. Mice were either treated with MTX (10 mg/kg of body weight, 3 times a week) or the vehicle control (VEH). MTX was dissolved at 2 mg/mL in a vehicle (VEH) containing 66% of PBS 1X and 33% of NaOH 0.1 M. MTX solution was filtered using a 0.22 μm filter and stored briefly in −20 °C until use.

Three times a week, starting the day of zymosan injection, mice were gavaged with MTX or VEH, and clinical arthritis scores (CAS) were evaluated. Experimenters evaluating CAS were blinded to the treatment. The wrist, ankle, and digits (4 digits on each upper limb and 5 digits on each lower limb) were scored. Joint swelling was monitored by inspection and scored as follows: 0, no joint swelling; 0.1,swelling of one finger joint; 0.5, mild swelling of wrist or ankle; 1.0, severe swelling of wrist or ankle. Scores for all fingers of forepaws and hindpaws, wrists and ankles were summed for each mouse. For example, an ankle that maintained its natural, angular shape was given a 0, an ankle with a round presentation was given a 0.5, and an ankle that presented a horseshoe shape due to severe swelling was given a 1. Similarly, if the wrist maintained its natural, slim appearance, it was given a 0, while some swelling was given a 0.5, and severe swelling that caused the wrist to bulge was given a 1. Each digit of fore paws and hind paws were scored as either 0 or 0.1 for no inflammation or inflammation, respectively. Each day, the score of each mouse’s digits, ankles, and wrists were added up and recorded as the CAS. The maximum CAS that a mouse could receive was 5.8 (1.5 for each lower limb plus 1.4 for each upper limb). CAS were collected over the course of the experiment and analyzed over time and stratified by microbiome or treatment. Additionally, individual AUCof CAS curves were calculated using the *auc* function from the MESS package (v0.5.12) in R.

#### 16S-seq of SKG mouse gut microbiota

Broadly, aliquots of mouse fecal samples were homogenized using a bead-beating (Mini-Beadbeater-24, BioSpec) method followed by DNA extraction and purification as previously described^47^. DNA was extracted using ZymoBIOMICS 96 well MagBead ZymoBIOMICS 96 MagBead DNA Kit (Cat#D4302) as per the manufacturer’s protocol. GoLay-barcoded 515F/806R primers^48^ were used to carry out 16S rRNA gene PCR according to the methods of the Earth Microbiome Project (earthmicrobiome.org). The following were combined: 2 µL of DNA, 25 µL of AmpliTaq Gold 360 Master Mix (Life Technologies), 5 µL of primers (2 µM each GoLay-barcoded 515/806R), and 18 µL H₂O. Amplification was as follows: 10 min at 95 °C, 25x (30 s at 95 °C, 30 s at 50 °C, 30 s at 72 °C), and 7 min at 72 °C. For the dose-response gnotobiotic experiment, amplicons were quantified with PicoGreen (Quant-It dsDNA; Life Technologies) and pooled at equimolar concentrations. Libraries were quantified (NEBNext Library Quantification Kit; New England Biolabs) and sequenced with a 600 cycle MiSeq Reagent Kit (251×151; Illumina) with ∼10% PhiX. For the remaining sequencing experiments, samples underwent primary PCR for amplification and secondary PCR to add flow cell adaptors and indices as previously described^49^. DNA was normalized to equal concentrations based on PicoGreen quantification and pooled. Pooled libraries were purified and concentrated with MinElute PCR Purification kit (Qiagen #28004), run on 1% gel, size-selected and purified using MinElute Gel Extraction kits (Qiagen, #28604). Pooled libraries were run at the Chan Zuckerberg Biohub using Illumina MiSeq platform.

### QUANTIFICATION AND STATISTICAL ANALYSIS

#### General

Power calculations were performed for PK and arthritis experiments whenever pilot data were available. All statistical analyses were performed using the R environment. “N.S.” indicates “not significant.” In general, Student’s t-tests were used to infer significance (p ≤ 0.05) unless otherwise indicated in the figure legend. Whenever provided, boxplots depict the median, hinges represent the first and third quartiles, and whiskers show the range. Whenever provided, error bars represent the standard error of the mean (SEM).

#### 16S rRNA amplicon analysis of mouse samples

Reads that were not already demultiplexed were demultiplexed using QIIME^50^ v1.9.1 (split_libraries_fastq.py). QIIME2^51^(v 2020.2) was used to trim reads, denoise the data, and create a feature table using the following commands: *qiime cutadapt trim-paired*, *qiime dada2 denoise-paired*, *qiime feature-table filter-samples*. Within QIIME2, taxonomy was assigned using the DADA2^52^ implementation of the RDP Naïve Bayesian classifier^53^ using the DADA2-formatted SILVA v128 training set; taxonomic assignments to the species level were chosen by exact match of the ASV to the reference database, and where multiple matches were present, all are reported. For figures and in the text, genus and species assignments are reported when available, and when neither was available, we included the family taxonomic assignment; we also report a unique ASV identifier. This ASV identifier can be used to look up the full taxonomic assignment from kingdom to species and the associated sequence variant. The QIIME2R package was used to import QIIME2 artefacts into R (https://github.com/jbisanz/qiime2R). Diversity metrics were generated using Vegan (v2.5-6) and Phyloseq (v1.30.0)^54^, with principal coordinate analysis (PCoA) or principal components analysis (PCA) carried out with Ape (v5.3) or Vegan, respectively. Analyses were carried out on either: (1) centered log2-ratio (clr) normalized taxonomic abundances calculated as A_clr_=[log_2_(A_1_/g_a_), log_2_(A_2_/g_a_),… log_2_(A_n_/g_a_),], where A is a vector of non-zero read counts and g_a_ is the geometric mean of all values of A, or (2) relative abundance calculated as proportion of reads. ANOSIM and PERMANOVA were used to detect changes in community composition using counts from rarefied data and Bray-Curtis distances. DESeq2 (v1.26.0)^55^ was used to determine differentially abundant taxa on raw count data. For each sample, fastq files are available in NCBI’s Sequence Read Archive (SRA), accession number [*to be determined upon publication*].

## Author Contributions

Conceptualization: RRN. Funding acquisition: JUS, PJT, RRN. Investigation: VSVS, ALD, CMH, DAO, ERR, MM, NP, SY, JA, RRN. Methodology: ALD, DAO, MM, ERR, ADP, JGB, DSD. Project administration: RRN. Resources: RBB, JUS. Supervision: JA, JUS, ADP, PJT, RRN. Validation: VSVS, ALD, CMH, RRN. Visualization: VSVS, ALD, CMH, DAO, RRN. Writing – original draft: RRN. Writing – review & editing: all authors.

## Acknowledgements

This work was supported by the following: A.D. was supported by a Postdoctoral Fellowship from the Belgian American Educational Foundation (B.A.E.F.); D.A.O. was supported by: 3K08AR073930-05S1 (NIAMS/OD) and ImmunoDiverse SRA Fellowship. R.B.B. was supported by NIH/NCATS UL1TR001445. E.R.R was supported by PSU/NIDDK (T32DK120509). This was work was supported by multiple grants: 2R01HL122593 (P.J.T), 1R01AR074500 (J.U.S and P.J.T), UCSF Breakthrough Program for Rheumatoid Arthritis-related Research (BPRAR; partially funded by the Sandler Foundation; P.J.T), Arthritis Foundation Center of Excellence (R.R.N); K08AR073930 (R.R.N); R03AR082036 (R.R.N.), I01CX002557 (R.R.N.), R35GM151349 (R.R.N.), Russell/Engleman Rheumatology Research Center (R.R.N.), and the Arthritis National Research Foundation (R.R.N.), Benioff Center for Microbiome Medicine (R.R.N. and P.J.T).

We acknowledge support from the following core facilities: UCSF QMAC, UCSF Gnotobiotics Core, PSU Metabolomics Core Facility at The Huck Institutes of the Life SciencesPenn State University (Sergei Koshkin).

We would like to thank William Seaman and members of the Nayak Lab for their helpful comments and suggestions on the manuscript.

---

**Supplementary Table 1.**
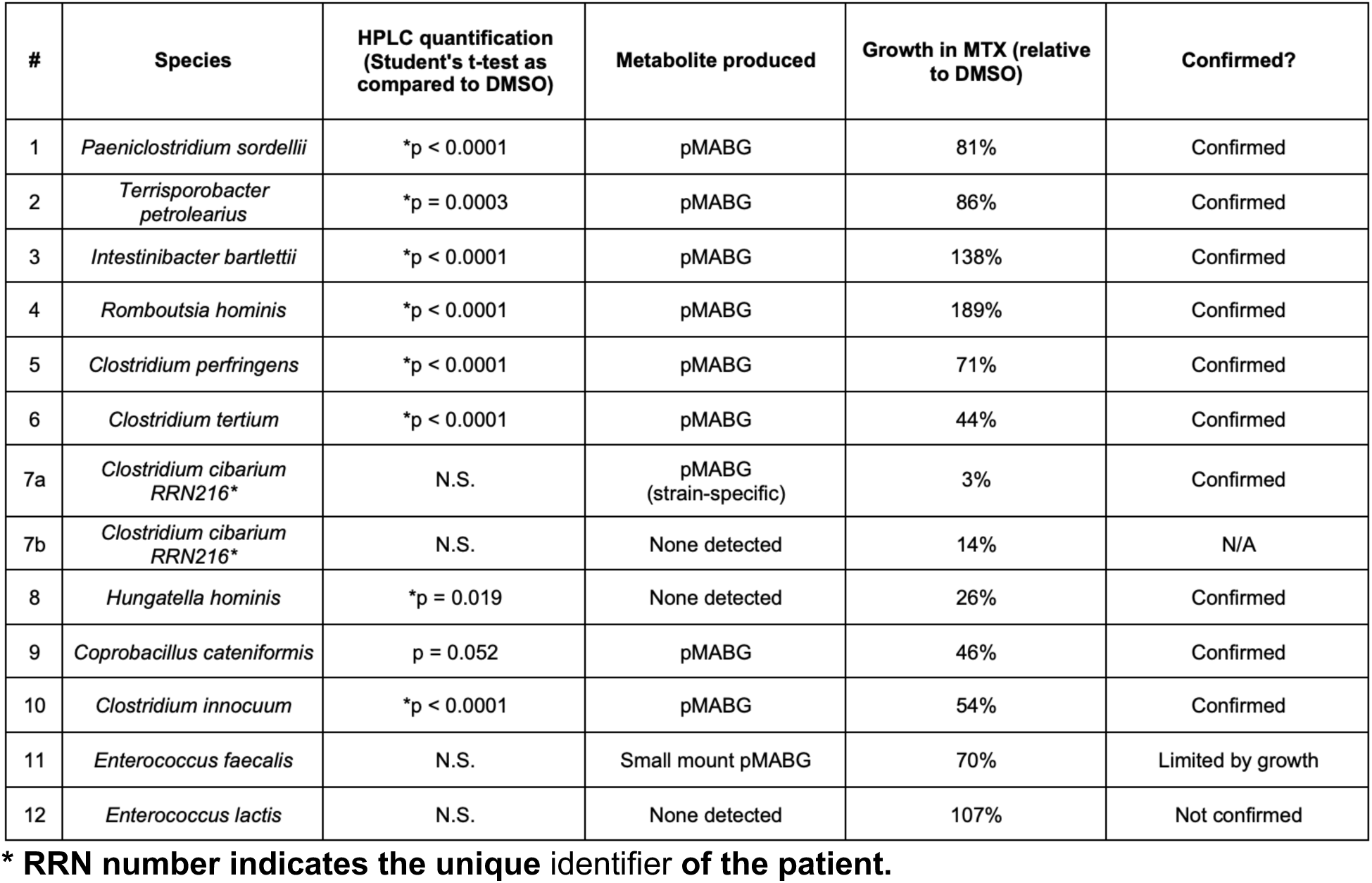
RA-associated human gut strains that metabolize MTX.

**Supplementary Table 2. Differentially abundant amplicon sequence variants (ASVs) in Zion vs. Parnassus mice (not co-housed).**

*See Excel sheet*.

**Supplementary Table 3.** Details of mouse experiments.

| # | Phenotype | Description | #mice/<br>group | Strain | Microbiome | Dose | Route | Sex | Age at<br>time of<br>treatment | Colonization<br>time prior to<br>treatment | Duration of<br>drug<br>treatment |
| --- | --- | --- | --- | --- | --- | --- | --- | --- | --- | --- | --- |
| 1 | PK | GF vs. SPF | 6<br>8 | C57BL/6J | GF<br>SPF | 50<br>mg/kg | PO | F | 6-7<br>weeks old | not relevant | 1 single dose<br>of MTX |
| 2 | PK | IP vs. PO | 3<br>3 | C57BL/6J | SPF | 50<br>mg/kg | PO<br>IP | M | 8<br>weeks old | not relevant | 1 single dose<br>of MTX |
| 3 | PK | Patient 170 vs.<br>Patient 317 | 11<br>11 | C57BL/6J | Humanized with Patient 170<br>Humanized with Patient 317 | 50<br>mg/kg | PO | M | 9-13<br>weeks old | 2 weeks | 1 single dose<br>of MTX |
| 4 | Arthritis<br>(#1) | vehicle (VEH)<br>vs. MTX,<br>co-housed Parn.<br>& Zion | 8<br>8 | BALB/c<br>SKG | SPF from PSB or MTZ<br>SPF from PSB or MTZ | (VEH)<br>10<br>mg/kg | PO | F | 10-13<br>weeks old | 2 weeks of<br>co-housing | MTX 3 times<br>a week |
| 5 | Arthritis<br>(#2) | Parn. origin vs<br>Zion origin<br>(not co-housed) | 6<br>3 | BALB/c<br>SKG | SPF from PSB<br>SPF from MTZ | 10<br>mg/kg | PO | F | 11-12<br>weeks old | not relevant<br>(no co-housing) | MTX 3 times<br>a week |
| 6 | PK | Parn. vs. Zion | 5<br>6 | BALB/c<br>SKG | GF colonized with Parn.<br>microbiota<br>GF colonized with Zion<br>microbiota | 50<br>mg/kg | PO | F | 13-24<br>weeks old | 2 weeks | 1 single dose<br>of MTX |

F: female; GF: germ-free, IP: intraperitoneal; M: male; MTX: methotrexate; Parn.: Parnassus; PO: per os (= oral); PK: pharmacokinetic; SPF: specific-pathogen-free; Zion: Mount Zion.

**Supplementary Table 4.** Details of LC-MS targeted metabolomics for *in vitro* and *in vivo* experiments.

| Compound | Q1 mass | Q3 mass | EP | CE | CXP | Mode | LC-MS/MS method |
| --- | --- | --- | --- | --- | --- | --- | --- |
| methotrexate (MTX) | 455.2 | 308.2 | 10 | 20 | 14 | Positive | A |
|  |  |  |  | 29 | 15 | Positive | B |
| methotrexate diglutamate (MTXPG2) | 584.2 | 308.2 | 10 | 20 | 14 | Positive | A |
|  |  |  |  | 36 | 15 | Positive | B |
| deoxyaminopteroic acid (DAMPA) | 326.2 | 175.1 | 10 | 20 | 14 | Positive | A |
|  |  |  |  | 26 | 15 | Positive | B |
| 7-hydroxy-methotrexate (7-OH-MTX) | 471.2 | 324.1 | 10 | 15 | 14 | Positive | A |
|  |  |  |  | 25 | 15 | Positive | B |
| 6-methylpteridine-2,4-diamine (6-MPDA) | 177.0 | 160.0 | 10 | 18 | 14 | Positive | A |
|  | 177.0 | 97.8 | 10 | 21 | 15 | Positive | B |
| aminopterin (APT, IS1) | 441.2 | 294.0 | 10 | 20 | 14 | Positive | A |
|  |  |  |  | 30 | 15 | Positive | B |
| 2-amino-3-bromo-5-methylbenzoic acid (2A3Br5MBA, IS2) | 231.0 | 104.0 | 10 | 25 | 15 | Positive | B |
| p-methylaminobenzoylglutamic acid (pMABG) | 278.9 | 150.2 | -10 | -25 | -15 | Negative | B |

CE: collision energy; CXP: collision cell exit potential; EP: entrance potential; IS: internal standard.

**Supplementary Figure 1.**
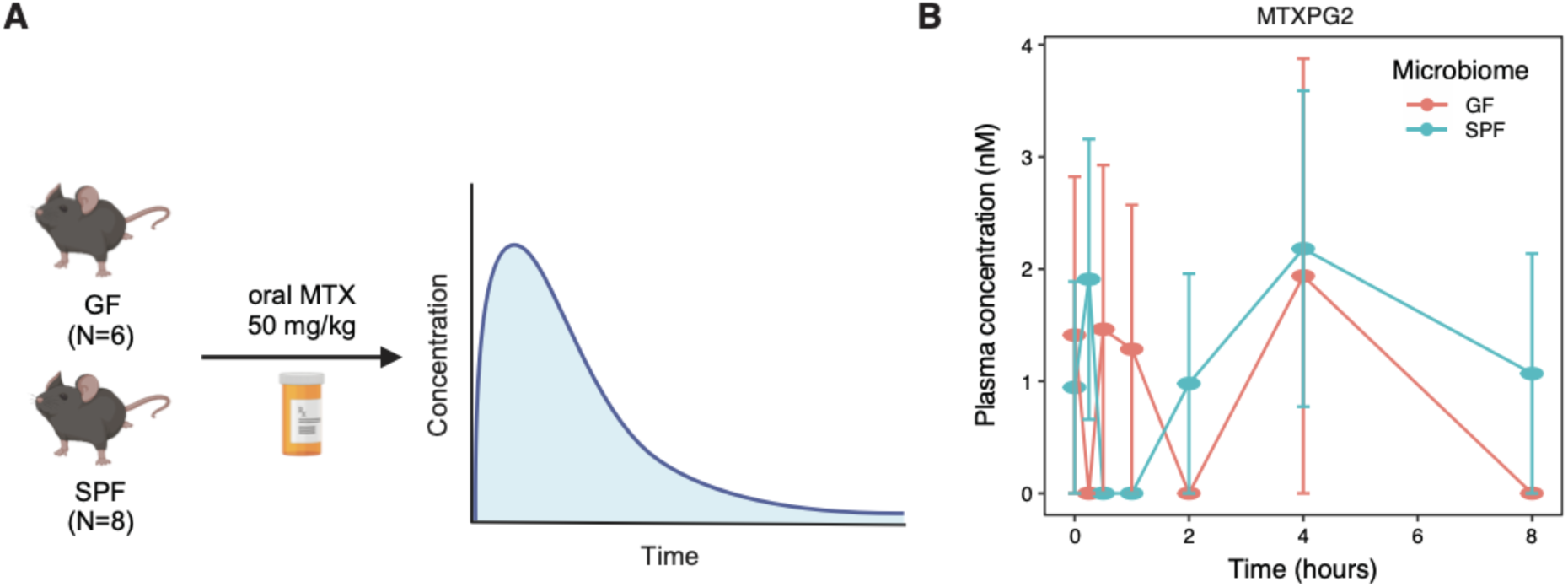
related to Figure 1. Experimental design and polyglutamated MTX. (**A**) Experimental design of a pharmacokinetic (PK) study with germ-free (GF, N=6) and colonized (SPF, N=8) mice. (**B**) MTX-PG_2_ levels in plasma.

**Supplementary Figure 2.**
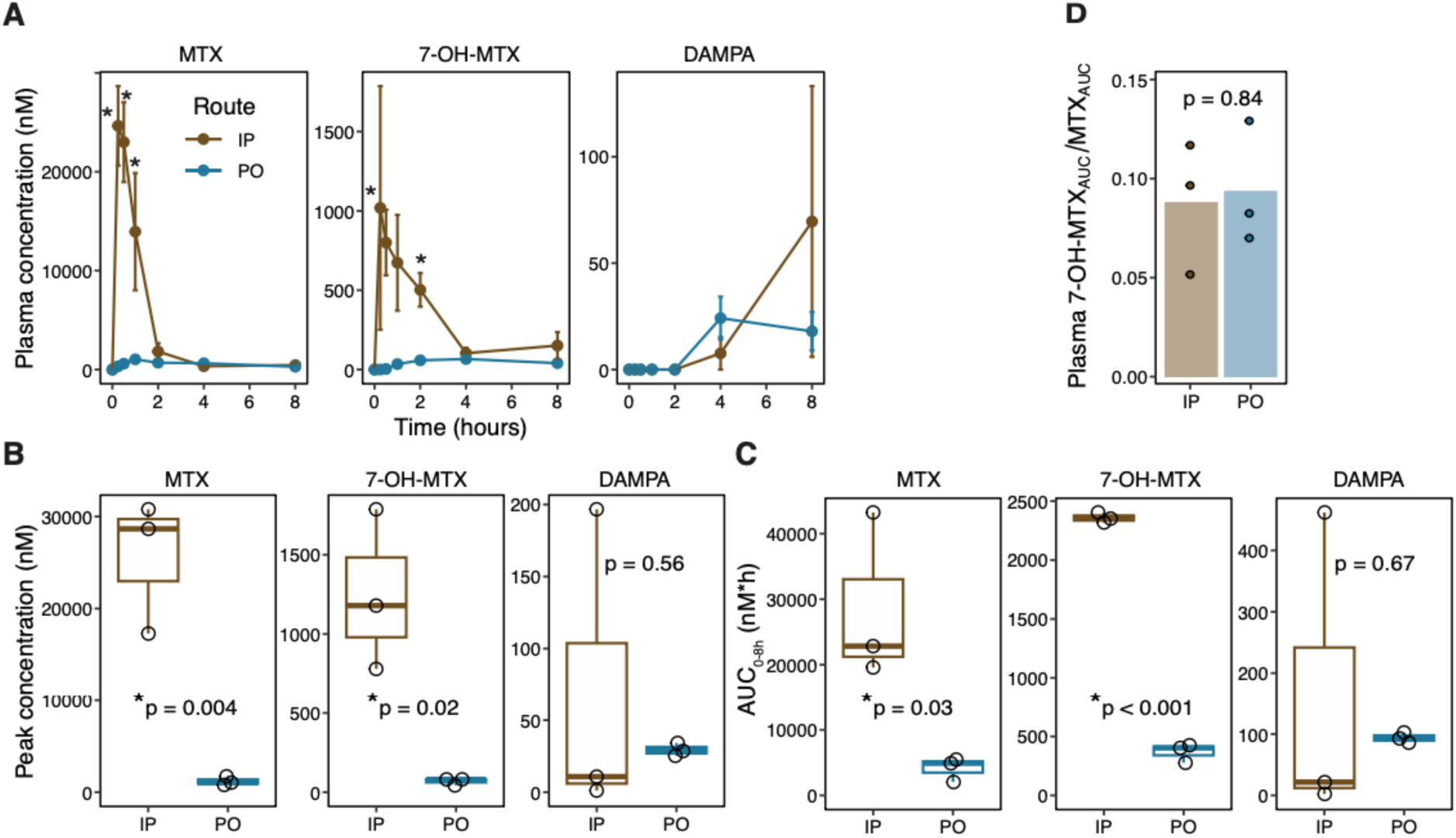
Parenteral administration of MTX does not preclude microbial metabolism of MTX into DAMPA. (**A**) Plasma concentration of MTX and its known metabolites were quantified by LC-MS in SPF C57BL/6J male mice after a one-time administration of MTX 50 mg/kg, given either by intraperitoneal injection (IP, N=3) or orally (PO, N=3). Multiple plasma samples were taken from each mouse over an 8 h time course. (**B**) Peak concentration and (**C**) AUC were compared between IP vs. PO treated mice. (**D**) Ratio of AUC of 7-OH-MTX to MTX in the plasma over 8 h. Results from Student’s t-test are reported. Error bars are SEM.

**Supplementary Figure 3.**
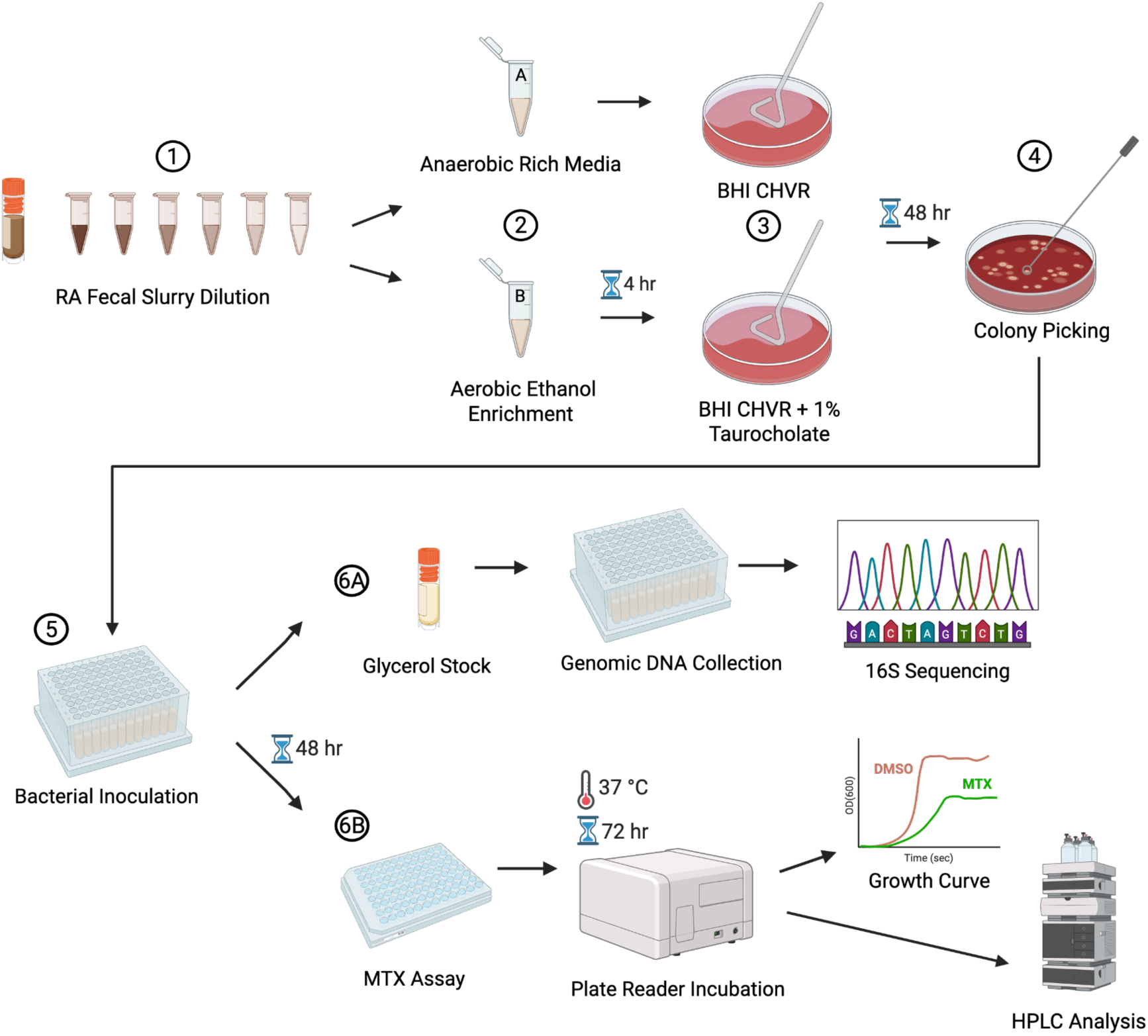
related to Figure 5. Strain harvesting workflow for identifying RA-associated microbial strains that metabolize MTX. (**Top panel, Steps 1 through 4**) Fecal slurries from RA patients are streaked out at multiple dilutes on rich media (BHI+), and media supplemented with taurocholate, a bile acid which induces spores to germinate. This latter step was devised to enrich for *Firmicutes*, which we found are enriched for MTX metabolizing species. (**Middle panel**) We picked 24 colonies with different morphologies per patient and created glycerol stocks. These glycerol stocks underwent 16S full-length Sanger sequencing in order to identify bacterial species (Step 5 and 6A). (**Bottom panel**) In Step 5, we also inoculated harvested isolates in either MTX 50 ug/mL or vehicle control (DMSO). We quantified growth to ensure that microbes grew at this concentration, incubating samples for 72 h. Cell pellets were spun down, and supernatant was extracted in methanol and profiled by HPLC to quantify the amount of MTX remaining compared to sterile controls (Step 6B). Figure generated with BioRender.

**Supplementary Figure 4.**
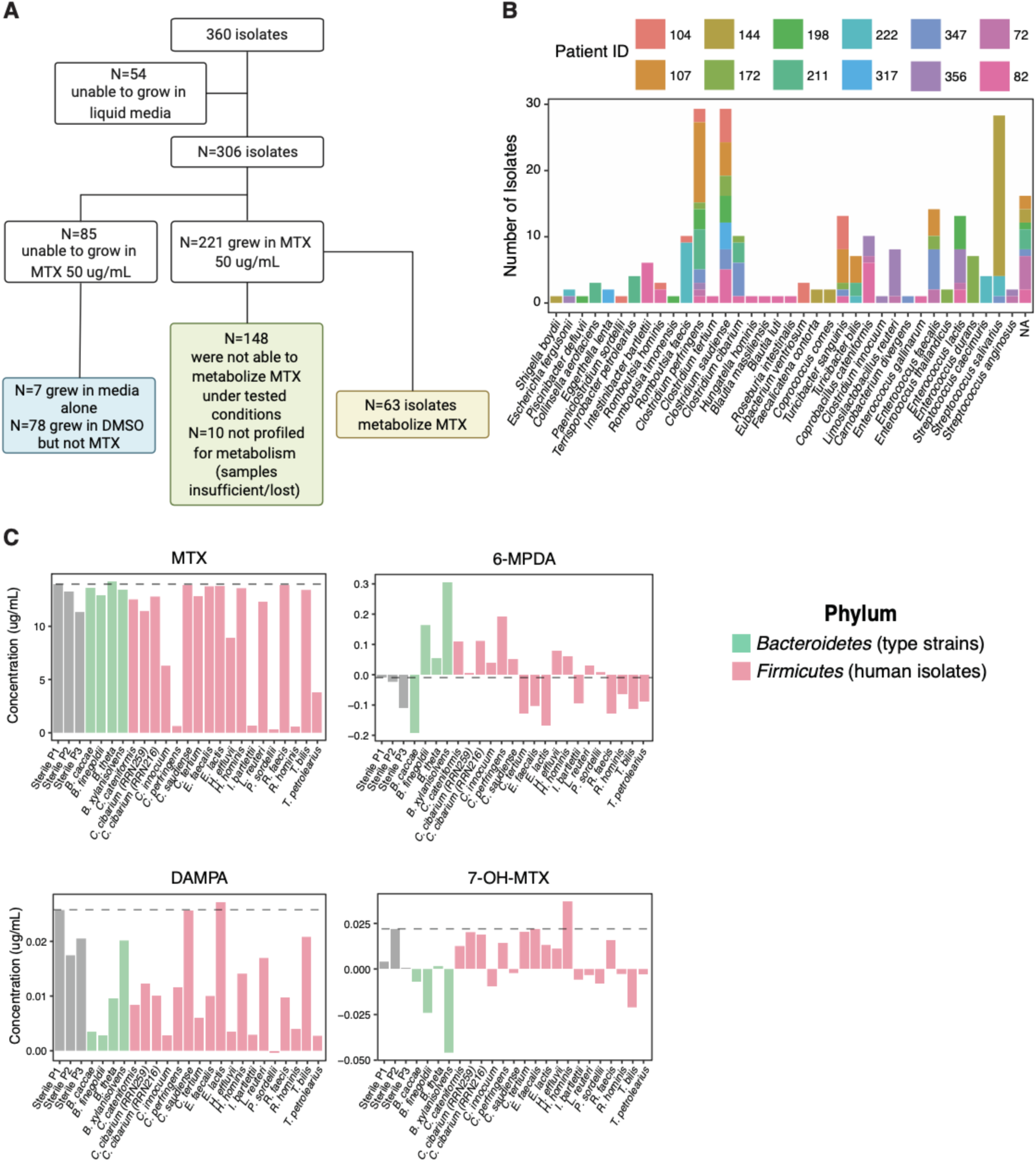
related to Figure 5. Many RA-associated bacterial species metabolize MTX into known and novel metabolites *in vitro*. **(A)** Flow chart of RA-associated microbes that were harvested and screened for MTX metabolism. (**B**) Unique species identified by full-length 16S Sanger sequencing. The contribution of each treatment-naïve RA patient donor is indicated by the color of the bar. (**C**) LC-MS quantification of MTX and its metabolites in the supernatant of RA-associated isolates incubated with MTX for 72 h (N=1 bacterial sample). The phylum is indicated by the color of the bar. Type strains of *Bacteroides* species (green) were used as negative controls.

**Supplementary Figure 5.**
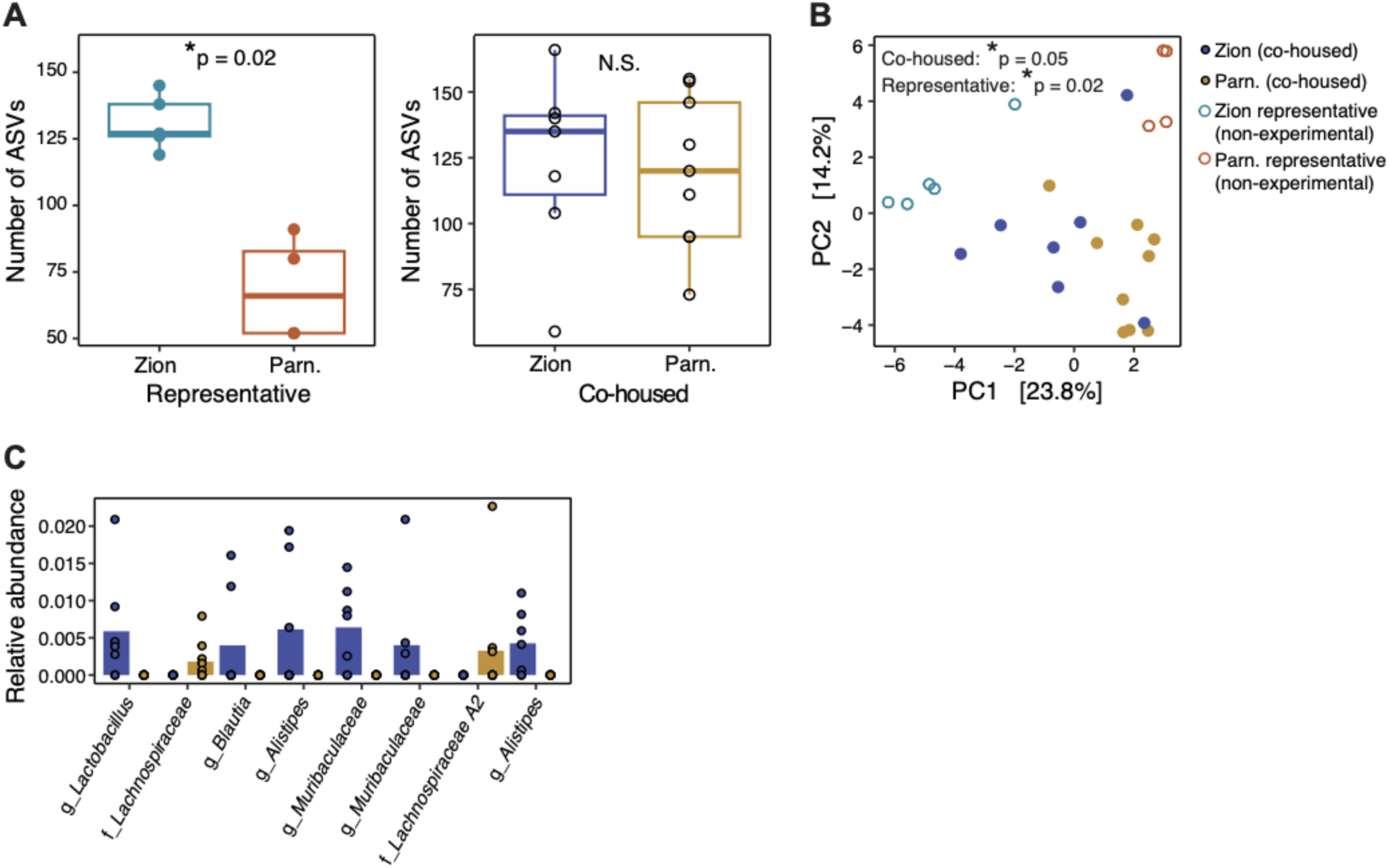
related to Figure 6. The microbiome of SKG mice reared at two different facilities is associated with MTX response. (**A**) Alpha diversity (number of observed ASVs) by facility-of-origin in mice that are not co-housed (“Representative”, left panel) and in mice that are co-housed for 2 weeks (“Co-housed”, right panel). (**B**) Principal components analysis of Euclidean distances of clr-transformed abundances (beta diversity) of Co-housed mice (solid circles) and Representative mice from each facility that were not co-housed (open circles). (**C**) Relative abundance of ASVs that were differentially abundant by facility-of-origin (DESeq, p_adj_<0.05) in mice that were co-housed. Results of PERMANOVA reported in B. All results in **C** are significant by DESeq.

